# Revealing the host antiviral protein ZAP-S as an inhibitor of SARS-CoV-2 programmed ribosomal frameshifting

**DOI:** 10.1101/2021.05.31.445667

**Authors:** Matthias M. Zimmer, Anuja Kibe, Ulfert Rand, Lukas Pekarek, Luka Cicin-Sain, Neva Caliskan

**Author notes:** These authors contributed equally to this work (Matthias M. Zimmer, Anuja Kibe).

## Abstract

Programmed ribosomal frameshifting (PRF) is a fundamental gene expression event in many viruses including SARS-CoV-2, which allows production of essential structural and replicative enzymes from an alternative reading frame. Despite the importance of PRF for the viral life cycle, it is still largely unknown how and to what extent cellular factors alter mechanical properties of frameshifting RNA molecules and thereby impact virulence. This prompted us to comprehensively dissect the interplay between the host proteome and the SARS-CoV-2 frameshift element. Here, we reveal that zinc-finger antiviral protein (ZAP-S) is a direct and specific regulator of PRF in SARS-CoV-2 infected cells. ZAP-S overexpression strongly impairs frameshifting and viral replication. Using *in vitro* ensemble and single-molecule techniques, we further demonstrate that ZAP-S directly interacts with the SARS-CoV-2 RNA and ribosomes and interferes with the folding of the frameshift RNA. Together these data illuminate ZAP-S as *de novo* host-encoded specific inhibitor of SARS-CoV-2 frameshifting and expand our understanding of RNA-based gene regulation.

## INTRODUCTION

The novel severe acute respiratory syndrome-related coronavirus (SARS-CoV-2), the causal agent of Coronavirus Disease 2019 (COVID-19), emerged rapidly to become a global threat to human health ^1^. Global analyses of RNA- and protein-interaction networks have increased our understanding of SARS-CoV-2 viral replication in a short time ^2,3^. However, there is a lack of detailed mechanistic understanding of the interplay between RNA-protein complexes, which could inform the design of novel antivirals. Here, functionally important RNA elements of the viral genome represent ideal targets due to their evolutionary conservation. One of those well-conserved RNA elements is the programmed ribosomal frameshift site.

A hallmark of infections by the SARS-CoV-2 and many other viruses is the –1 programmed ribosomal frameshifting (–1PRF) event which allows translation of multiple proteins from the same transcript. Frameshifting increases the coding potential of the genomes and is often used to expand the variability of proteomes, adapt to changing environments, or ensure a defined stoichiometry of protein products ^4,5^. In coronaviruses, –1 frameshifting on the *1a*/*1b* gene is fundamental for efficient viral replication and transcription of the viral genome. In cells, efficiency of this frameshifting event varies between 20-40% ^6,7^. Programmed ribosomal frameshifting relies on the presence of a slippery heptameric sequence (in coronaviruses U UUA AAC) and an RNA secondary structure such as a pseudoknot **(Fig. 1A)**. Mutations in the slippery sequence and downstream RNA structure drastically impair frameshifting efficiency ^8,9^

**Fig. 1.**
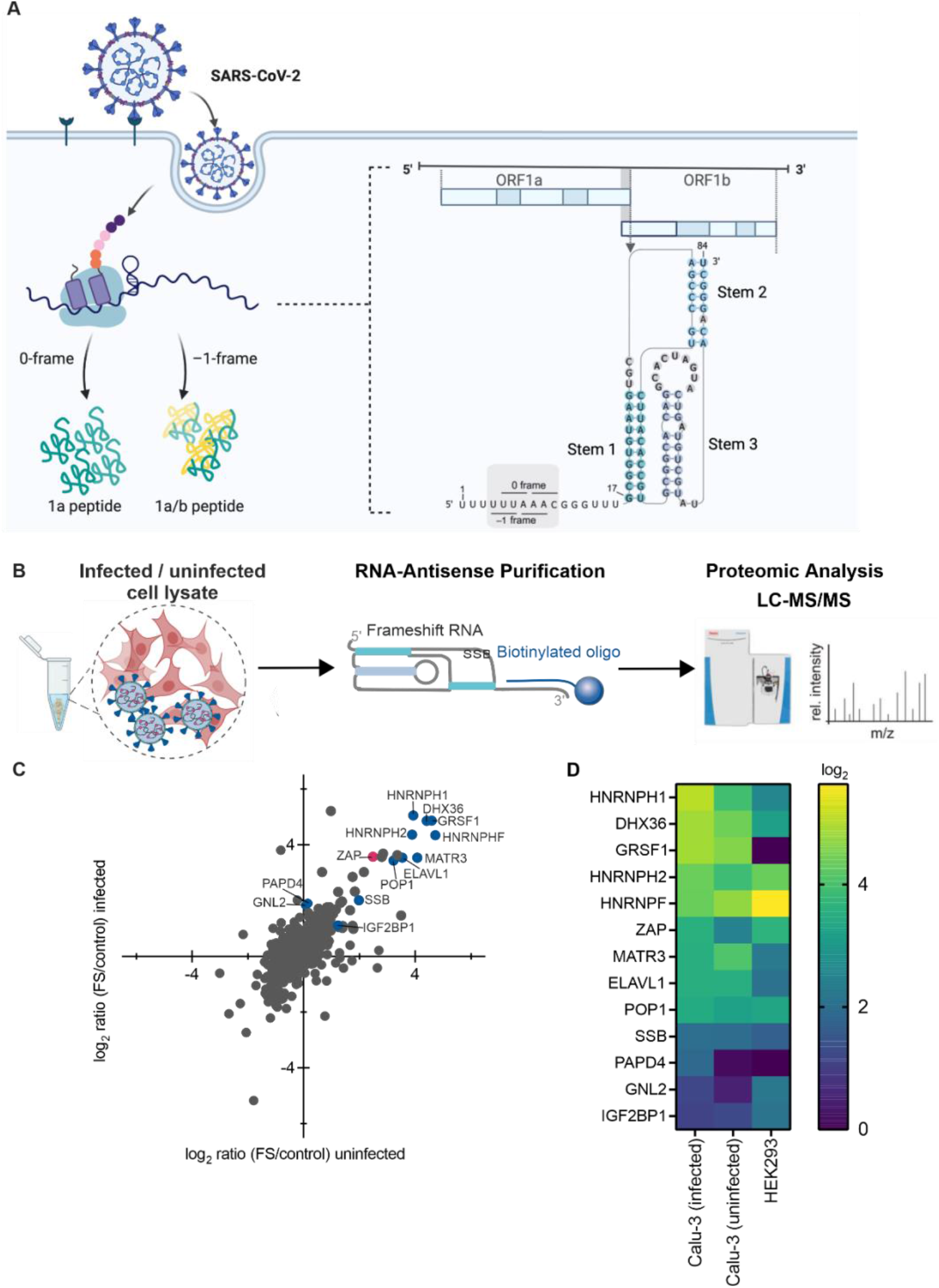
*In vitro* RNA-antisense purification-based discovery of protein interactors of the SARS-CoV-2 –1PRF element. **(A)** Schematic representation of the relevant genomic segment of SARS-CoV-2 as well as the location of the –1PRF element. **(B)** Schematic of *in vitro* interactome capture of protein interactors of the SARS-CoV-2 –1PRF element. *In vitro* synthesized RNA fragments corresponding to nucleotides 13456-13570 of the SARS-CoV-2 genome were incubated with lysates of naïve HEK293 cells as well as SARS-CoV-2-infected and uninfected Calu-3 cells. The –1PRF RNA was captured by a biotinylated antisense DNA oligo and isolated proteins were subjected to LC-MS/MS. **(C)** Representative scatter plot of log_2_-ratios comparing proteins captured in uninfected vs. SARS-CoV-2-infected Calu-3 cells. Core interactors common between uninfected and SARS-CoV-2-infected Calu-3 cells as well as uninfected HEK293 cells are highlighted in blue, ZAP is highlighted in pink. **(D)** Heat map representing the respective enrichment (log_2_) of core interactors. See also **Supplementary Fig. 1D and Supplementary Table 1**.

Traditionally, efforts to understand the mechanism of –1PRF focused on *cis*–acting modulatory elements. Previous work in purified translation systems explained in unprecedented detail how ribosome pausing on the slippery codons may lead to a kinetic partitioning and favor movement of translating ribosomes to an alternative reading frame ^6,10^. It has been shown that –1PRF may occur during a late stage of the tRNA translocation step with the stimulatory element causing ribosomes to become trapped in an unusual conformation that is relieved by either the spontaneous unfolding of the blockade or a –1 slip on the mRNA ^6,10^. Recently, it is becoming clear that *cis*-acting elements are not the only determinants of frameshifting in cells and *trans-acting* viral and cellular factors as well as small molecules or oligonucleotides can alter frameshifting levels ^11–13^. Despite this momentum, fundamental questions such as how pertinent RNA-binding factors are for frameshifting processes in general and how exactly these interactions alter the mechanical properties of RNA as well as the choice of the reading frame remain to be exploited.

Based on current knowledge, there would be at least three potential routes to modulate frameshifting by *trans*-acting factors. First, the binding of the factor can transform the downstream RNA element to a more stable roadblock, which was shown for cardiovirus 2A, poly-(C) binding protein and some small molecules such as the NCT-8 ^11,12,14^. In these cases, the specific interaction of the factor with the nucleotides downstream of the slippery codons leads to an increase in frameshifting. Alternatively, eukaryotic release factors such as eRF1 alone or eRF1/3 recruited by Shiftless (SFL) to the HIV-1 frameshift site were shown to target stalled ribosomes ^15,16^. In this case, different from the first group of regulators the interaction of both SFL and release factors was not dependent on the identity of the frameshift RNA. Therefore, it remains to be solved how the frameshifting ribosome complexes would be recognized by these *trans*-acting factors. A third route could potentially work through remodeling or destabilization of the frameshifting RNA elements through direct interactions between the RNA and the *trans*-factor. However, so far there has been no cellular or viral factor reported to affect frameshifting efficiency (FE) through this route.

These prompted us to comprehensively identify and study direct interactions between the host cell proteome and the SARS-CoV-2 frameshifting RNA element. Firstly, to decipher interactors of the frameshifting RNA element, we employed an *in vitro* RNA-antisense capture and mass spectrometry-based screen ^17^. Through this approach, we identified the short isoform of zinc-finger antiviral protein (ZAP-S, ZC3HAV1), as a prominent RNA interaction partner. We demonstrated that ZAP-S acts as a host-encoded inhibitor of SARS-CoV-2 1a/1b frameshifting *in vivo* and *in vitro*. Intriguingly, ZAP-S overexpression reduced the replication of SARS-CoV-2 by more than 90%, highlighting the importance of the protein in the viral life cycle. The effect of ZAP-S on SARS-CoV-2 frameshifting was specific, because barring the closely related SARS-CoV-1, other viral and cellular PRF levels were not affected by ZAP-S *in vivo*. Using a multidisciplinary approach, we further probed this effect and revealed important clues on molecular principles of frameshifting downregulation by ZAP-S. Amongst them, we show that ZAP-S can alter the physical properties of the PRF RNA, which brings a unique dimension to frameshift mechanisms. Our study highlights for the first time that the expression of the SARS coronavirus ORF1a/1b, can be directly and specifically modulated by a host-encoded RNA-binding protein during infection. These findings provide substantial new insights on PRF regulation and the interplay between SARS-CoV-2 replication and host defense, thereby paving the way for novel RNA-based therapeutic intervention strategies.

## RESULTS

### SARS-CoV-2 PRF RNA capture identifies novel host interactors

To identify potential cellular RNA-binding proteins (RBPs) that interact with the –1PRF element of SARS-CoV-2, an *in vitro* synthesized RNA fragment corresponding to nucleotides 13456-13570 of the SARS-CoV-2 genome was incubated with lysates of SARS-CoV-2-infected and uninfected Calu-3 cells and naïve HEK293 cells **(Fig. 1B)** ^17^. Calu-3 cells are lung epithelial cells that are commonly used to study CoV infection ^18^. HEK293 cells are routinely used to study RNA-protein interactomes, therefore they represented an ideal system to assess possible cell-based variations ^19^. To exclude any non-specific binders, we used an 80 nucleotides long non-structured RNA as a control. RNAs were captured by a biotinylated antisense DNA-oligo, and interacting proteins were identified by LC-MS/MS (liquid chromatography tandem mass spectrometry) analysis **(Fig. 1B, C)**.

In our SARS-CoV-2 frameshift RNA capture, more than 100 proteins were at least two-fold enriched. According to our GO term analysis, the majority (80%) of identified hits have been described as RNA-binding proteins **(Supplementary Fig. 1A)**. As for viral proteins, we saw an enrichment of the viral nucleocapsid protein (N) in infected lysates, which is a well-described RNA-binding protein ^20^. In addition, 35% and 30% of the enriched RBPs were involved in splicing and ribosome biogenesis, respectively **(Supplementary Fig. 1A)**. Among those, 19 proteins were common to infected and uninfected Calu-3 cells, 18 hits were identified only in HEK293 cells, 15 were captured only in uninfected Calu-3 cells, and 40 were present only in infected Calu-3 cells **(Supplementary Fig. 1B)**. The core interactome of 9 proteins identified in all three cell systems encompasses well-described post-transcriptional regulators **(Fig. 1C, D, Supplementary Fig. 1C, Supplementary Table 1)**. Proteins recently identified in genome-wide interactome studies as direct RNA interaction partners for SARS-CoV-2 were selected for downstream functional characterization ^3,20–22^. Several of these have been shown to play a role in RNA processing, including splicing (such as HNRNPs F, H1, and H2), RNA trimming (POP1) and RNA decay (ZAP) ^23–25^. It is well known that some nuclear RBPs can localize and may carry out diverse functions to regulate gene expression in the cytoplasm ^23^. For instance, HNRNP A2/B1 binds to VEGF-A mRNA and thereby affects recoding through readthrough ^26^, and HNRNPD is involved in RNA folding and replication of Flaviviruses ^27^. Translational regulators included IGF2BP1, ELAVL1, DHX36, and SSB ^28,29^. ELAVL1 is a cofactor which ensures translational fidelity in the context of uORFs ^30^. DHX36 functions as a multifunctional helicase and is involved in translation and innate immunity ^31,32^. ZAP is an antiviral protein with two isoforms (ZAP-S and ZAP-L), both of which are implied in various RNA-related mechanisms, including RNA decay and translation ^25,33^. In addition to the aforementioned proteins, three more hits were selected for further analysis based on their fold enrichment in infected Calu-3 lysates. These included the RNA-binding protein GRSF1, the poly(A) polymerase PAPD4 as well as GNL2 which has been implied in ribosome biogenesis ^34–36^. Unlike IGF2BP1, the closely related IGF2BP3 was enriched to a much lesser extent (log_2_ enrichment 0.4-0.7) and, hence, was included as a control for the functional assays.

### RNA interactors specifically inhibit SARS-CoV-2 frameshifting in cells

In order to explore the potential role of the RNA binders in SARS-CoV-2 frameshifting, we designed an *in vivo* fluorescence-based –1PRF assay. In this assay, the expression of the first ORF EGFP in the 0-frame would be constitutive, whereas the expression of the following ORF mCherry would depend on – 1PRF occurring at the preceding SARS-CoV-2 1a/1b frameshifting fragment **(Fig. 2A)**. As controls, we used a construct lacking the –1PRF stimulatory sequence, and the mCherry gene is placed either in –1 or in-frame with respect to EGFP **(Fig. 2A, B)**. Frameshift efficiencies were calculated as the ratio of mCherry to EGFP in the test construct normalized to the in-frame control **(see also Materials and Methods)**. To study the effect of the *trans*-acting factors on SARS-CoV-2 1a/1b frameshifting, cells were co-transfected with both the dual-fluorescence reporter plasmid and the plasmid encoding the putative *trans*-factor as an N-terminal ECFP fusion. This was advantageous over luciferase-based assays, because fluorescence readout allowed gating for the ECFP+ cells, which express the protein of interest (see also **Supplementary Fig. 2F**). To benchmark the assay, a vector expressing only ECFP was used as a control to compensate for the spectral overlap between ECFP and EGFP. In addition, vector expressing ECFP-SFL, a previously described inhibitor of –1PRF in SARS-CoV-2, was used as a positive control ^3^. Using this fluorescence reporter system, the frameshifting efficiency (FE) of SARS-CoV-2 was measured to be ca. 35% in HEK293 **(Fig. 2C, D, Supplementary Fig. 2A, B, Supplementary Table 2)**, which agreed well with the published FE for SARS-CoV-1 as well as the SARS-CoV-2 ^7,9,37^. As expected, deletion of the nucleotides corresponding to the predicted stem 2 of the pseudoknot as well as deletion of the region corresponding to the predicted stem 2 and 3 led to the abrogation of –1PRF, in line with minimal sequence requirements for frameshifting of other coronaviruses **(Supplementary Fig. 2A)** ^9^.

**Fig. 2.**
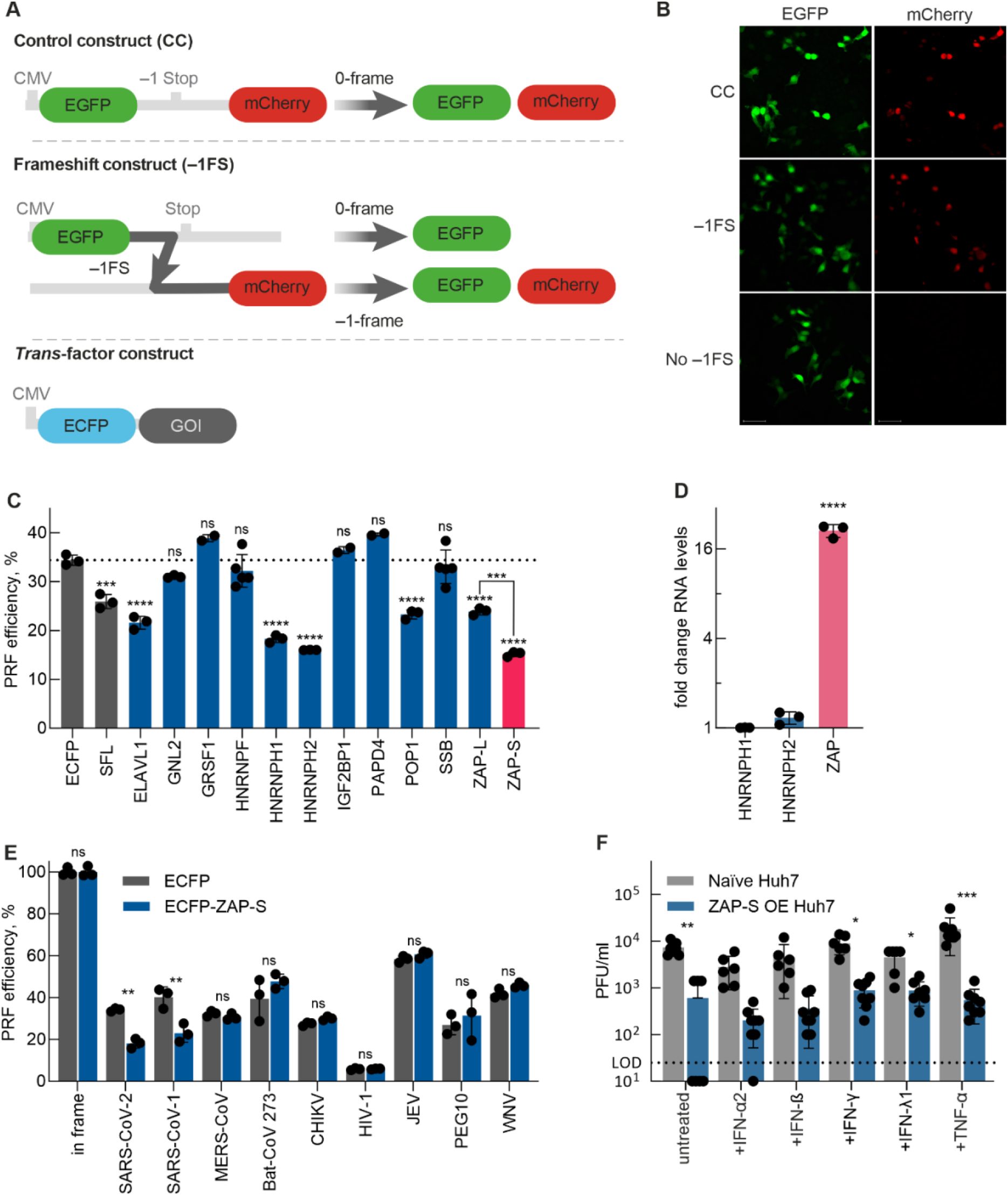
A functional screen of SARS-CoV-2 –1PRF element interactors. **(A)** Schematic representation of the dual-fluorescence frameshift reporter construct. The control construct consists of EGFP in-frame with mCherry separated by a self-cleaving 2A peptide resulting in equimolar production of EGFP and mCherry. In the frameshift construct below, EGFP and mCherry are not only separated by a self-cleaving 2A peptide but also by a stop codon in-frame with EGFP. As a result, 0-frame translation would produce only EGFP, whereas –1PRF would produce both EGFP and mCherry. The ratio of mCherry to GFP fluorescence is used to quantify the FE. The *trans*-factor construct is an N-terminal fusion of ECFP with the protein of interest to be analyzed. The control construct consists of ECFP alone. **(B)** Confocal microscopy images of cells transfected with EGFP-mCherry control (CC), –1FS, and no FS (no –1PRF site included after EGFP and mCherry in-frame with EGFP) constructs. The size bar represents 50 μm. **(C)** Comparison of relative FE of cells overexpressing *trans*-factors as ECFP fusion proteins. Data points represent the mean ± s.d. (n = 3 independent experiments). P values were calculated using an ordinary unpaired one-sided ANOVA comparing every condition to the ECFP control. ZAP-L and ZAP-S were separately compared to each other. * P < 0.01 – ** P < 0.001 – *** P < 0.0001 – **** P < 0.00001. **(D)** Expression profiles of selected genes in Calu-3 infected cells with SARS-CoV-2 at 72 hours. RNA levels were quantified by qRT-PCR and normalized to the respective RNA abundance in uninfected cells (shown as 2^-ΔΔCT^). Data points represent the mean ± s.d. (n = 3 independent experiments). P values were calculated using an ordinary unpaired one-sided ANOVA comparing delta CT values of the respective RNA in uninfected and infected cells. * P < 0.01 – ** P < 0.001 – *** P < 0.0001 – **** P < 0.00001. See also **Supplementary Fig, 2 and Supplementary Table 2**. **(E)** *In vivo* dual-fluorescence of additional –1PRF RNAs in HEK293 cells in the presence and absence of ZAP-S. SARS-CoV-1 – Severe acute respiratory syndrome-related Coronavirus 1, MERS-CoV – Middle East respiratory syndrome-related Coronavirus, Bat-CoV-273 – Bat Coronavirus 273, CHIKV – Chikungunya Virus, HIV-1 – Human Immunodeficiency Virus 1, JEV – Japanese Encephalitis Virus, WNV – West Nile Virus. Data points represent the mean ± s.d. (n = 3 independent experiments). P values were calculated using an ordinary unpaired one-sided ANOVA comparing every condition to the ECFP control. * P < 0.01 – ** P < 0.001. **(F)** Virus titers in the supernatant of infected naïve Huh7 or ZAP-S overexpressing Huh7 cells (ZAP-S OE) at 24 hours post infection. Treatment with IFN-γ (500 U/ml), IFN-β (500 U/ml), or IFN-λ1 (5 ng/ml) was done one hour before infection. Boxes show mean values ± s.d. (n = 4). The dotted line represents the limit of detection (LOD). ND: not detected.

Among the candidate PRF regulators examined, we observed no change in FE with GNL2, HNRNPF, IGF2BP1 or SSB which points to the fact that binding to the stimulatory RNA element is not sufficient for modulating PRF. Also, several proteins that were not significantly enriched in the interactome capture were not able to lead to significant changes in FE corroborating the specificity of the flow-cytometry-based frameshifting assay **(Supplementary Fig. 2B)**. On the other hand, several candidates such as SFL, HNRNPH1, HNRNPH2, ZAP-L and ZAP-S, showed a substantial reduction in FE by up to 50% **(Fig. 2C)**. Other candidates like GRSF1 and PAPD4 led to a slight but not statistically significant increase in FE. The candidates with the strongest effect on FE were HNRNPH1, HNRNPH2 and ZAP-S. In order to test whether these RBPs were functionally relevant, we measured their relative expression levels using quantitative RT-PCR in SARS-CoV-2 uninfected versus infected Calu-3 cells at 72 hours post-infection. Only ZAP showed a significant ca. 20-fold increase of mRNA levels upon infection **(Fig. 2D)**.

ZAP is a well-known multi-functional antiviral protein with an established role in the innate immune response against alphaviruses, filoviruses, HIV-1, hepatitis B virus and influenza virus ^38^. Whilst the mechanism of action for ZAP seems to revolve around the modulation of RNA stability ^39–41^, a more recent study implies its role in the maturation of viral particles ^42^. The *ZAP* gene produces two isoforms, ZAP-S and ZAP-L, differing only in their N-terminal domains. The two ZAP isoforms were suggested to carry out different functions in infections ^33^. While the longer, constitutively expressed isoform ZAP-L preferentially targets viral RNAs, the shorter isoform ZAP-S has been identified as an immune-regulatory protein that targets interferon mRNAs ^33^. In our assay, both isoforms were tested simultaneously, and we observed the decrease in frameshifting upon overexpression of the ZAP-S and ZAP-L. In both cases, the level of 0-frame product EGFP was not changed **(Supplementary Table 2)**. However, the –1PRF-inhibitory effect of ZAP-S was significantly stronger than the one by ZAP-L **(Fig. 2C)**. Therefore, we decided to follow up the interactions of ZAP-S as a frameshift regulator in detail.

First, to further corroborate the specificity of ZAP-S for the SARS-CoV-2 frameshift element, we tested whether the overexpression of ZAP-S affects –1PRF of other mRNAs, e.g., different Betacoronaviruses (SARS-CoV-1, MERS-CoV, Bat-CoV-273), Arboviruses (West Nile Virus (WNV), Japanese Encephalitis Virus (JEV), Chikungunya Virus (CHIKV)), and Human Immunodeficiency Virus-1 (HIV-1). Our analysis also included the embryonic gene *PEG10*, which represents an established example for –1PRF in humans ^43^. ZAP-S overexpression led to inhibition of –1PRF for SARS-CoV-1 *in vivo* by 50%. Among the other frameshift sites we tested, including *PEG10*, none were responsive to ZAP-S **(Fig. 2E)**. Given the high degree of similarity between the SARS-CoV-1 and CoV-2 frameshift sites, our results demonstrate that the effect of ZAP-S on frameshifting of SARS-CoV is specific. This is unlike the SFL protein, which affects several PRF genes, including the cellular PEG10 ^3,15^.

### Impaired viral replication by ZAP-S

Next, to test that ZAP-S is relevant in the context of infection, Huh7 cells stably overexpressing the ALFA-tagged ZAP-S were infected with SARS-CoV-2. We were able to demonstrate that the viral replication was reduced by more than 90% in cells overexpressing ZAP-S after 24 hours, which was in line with previous reports ^20,44^. Confirming our results, recent literature also indicated that siRNA-mediated depletion of ZAP leads to an increase in viral replication ^20,44^. Conversely, in our experiments the addition of interferons IFN-α2, INF-ß. IFN-γ, and IFN-λ1 as well as TFN-α did not lead to an inhibition of SARS-CoV-2 replication alone or in combination with ZAP-S overexpression **(Fig. 2F, Supplementary Fig. 2C)** ^44^.

Using mutated frameshift elements, it has been reported that even 10% change in the –1 frameshift product compared to the in-frame protein product inhibited SARS-CoV viral propagation and reduced infectivity ^45^. Furthermore, protein-mediated inhibition of PRF was shown to lead to similar effects on viral infectivity ^14, 46^. Since ZAP-S levels are upregulated upon infection **(Fig. 2D)** ^20,44^, it is likely that the reduction in viral titers was at least partially due to impaired frameshifting and reduced the –1 frame produced RdRP levels. This is supported by the finding that ZAP-S has a stronger antiviral effect than ZAP-L on SARS-CoV-2 replication ^20^. To test that we attempted to quantify the amounts of the frameshift product RdRP (nsp12) in the infected lysates via western blotting and proteomics. However, due to the low sensitivity of the antibody for western blot, and different charges of the frameshift versus non-frameshift products for the MS, so far, we were not able to confidently quantify the level of frameshifted product in infected cells.

### ZAP-S decreases SARS-CoV-2 frameshifting efficiency *in vitro*

We next focused on characterizing ZAP-S mediated regulation of frameshifting *in vitro* to dissect the molecular basis for inhibition of –1PRF. In principle, ZAP-S can either directly interact with the frameshift site or form a complex with auxiliary proteins to modulate translation. To test that, we recombinantly expressed and titrated ZAP-S in the rabbit reticulocyte lysate (RRL) translation system. We employed reporter mRNAs containing nucleotides 12686-14190 of the SARS-CoV-2 genome to best mimic the native genomic context of viral frameshifting. Control RNAs exclusively producing either the 0-frame (nsp9-11) or –1-frame products (nsp9-11 + partial nsp12) were employed as size markers for the western blot **(Fig. 3A and 3B)**. In the absence of ZAP-S, SARS-CoV-2 FE was about 46% **(Fig. 3B)**, as has been previously reported ^7^. With increasing amounts of ZAP-S, there was a corresponding decrease in FE. At the highest concentration of ZAP-S (3 μM), FE was reduced to ~26% **(Fig. 3B and 3C, Supplementary Table 3)**. To confirm that the observed effect was mediated by ZAP-S and not by random RNA-protein interactions we also analyzed the change in frameshifting upon addition of IGF2BP3 and SUMO-tag alone. IGF2BP3 was identified in our RNA interactome screen, but it was not significantly enriched unlike its close relative IGF2BP1 (log_2_ enrichment >1.5), thus rendering it suitable as a control. Here, neither the addition of IGF2BP3, nor the addition of SUMO alone led to a change frameshifting levels **(Supplementary Fig. 2D, E)**.

**Fig. 3.**
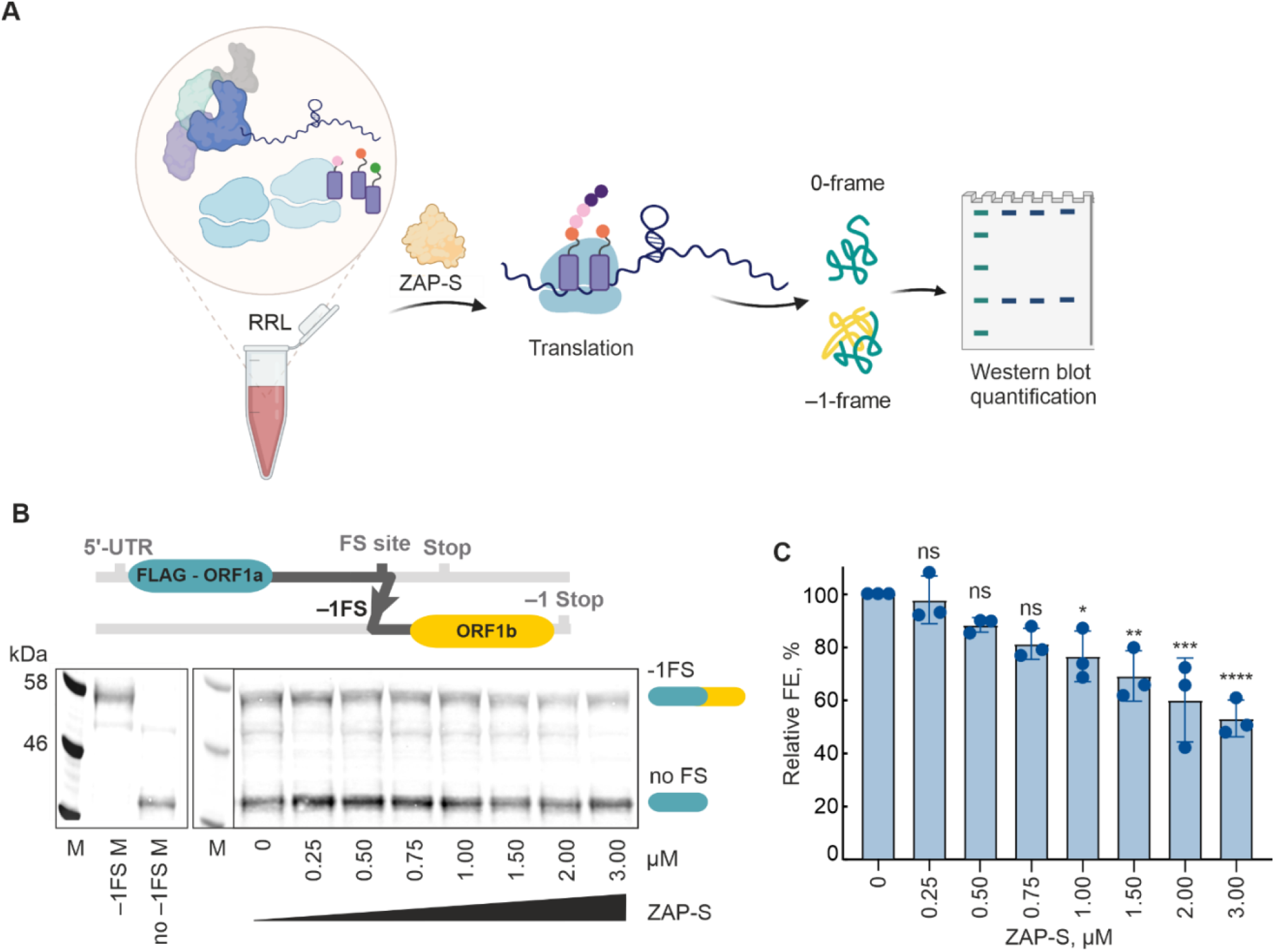
Effect of zinc-finger antiviral protein (ZAP) on 1a/1b –1 frameshifting *in vitro*. **(A)** The strategy of the *in vitro* translation assay using rabbit reticulocyte lysate (RRL). **(B)** Schematics of the N-terminal FLAG-tagged frameshifting reporter consisting of the nucleotides 12686-14190 (~1.5kb) of the SARS-CoV-2 genome. RNAs were translated in RRL in the presence of increasing concentrations of ZAP-S ranging from 0 to 3 μM. FLAG-tagged peptides generated by ribosomes that do not frameshift (no –1PRF) or that enter the –1 reading frame (−1PRF) were identified via western blotting using anti-DDDDK antibody. FE was calculated as previously described ^11^, by the formula: Intensity (–1-frame)/ (Intensity (–1-frame) + Intensity (0-frame)). Size markers - M (Marker), –1PRF M (–1 frame marker), and no –1PRF M (0-frame marker). **(C)** Changes in FE observed in the presence of ZAP-S from (B) (normalized to 0 μM ZAP as shown in B). P values were calculated using an ordinary unpaired one-sided ANOVA comparing every concentration to the no ZAP control. * P < 0.01 – ** P < 0.001 – *** P < 0.0001 – **** P < 0.00001. See also **Supplementary Fig. 2 and Supplementary Table 6**.

### ZAP-S directly interacts with the SARS-CoV-2 frameshifting motif

In order to dissect the interplay between SARS-CoV-2 frameshift stimulatory RNA structure and ZAP-S in detail, we prepared a series of short synthetic RNAs and we performed RNA-protein binding assays using the highly-sensitive microscale thermophoresis assay (MST). For that, wild-type (WT) RNA containing the –1PRF signal was derived from nucleotides 13456-13570 of the SARS-CoV-2 genome. The PRF RNA fragment was *in vitro* transcribed and Cy5-labeled at the 3’ end to monitor the change in fluorescence in response to ZAP binding.

With the wild-type SARS-CoV-2 pseudoknot, we observed that ZAP-S interaction occurs with the RNA with a high nanomolar affinity (K_D_ = 121 ± 12 nM) **(Fig. 4A and Supplementary Fig. 3A)**, which argued that ZAP-S is a direct interaction partner of the frameshift signal. We next employed mutant RNAs to detail the molecular basis of RNA: ZAP-S interaction. The first mutant, SL2 mutant, had a truncation at the 3’ end of the putative SARS-CoV-2 pseudoknot, which is predicted to prevent the formation of the putative stem 2 ^47^ **(Fig. 1A, Fig 4B)**. Furthermore, the bulged adenine residue in stem 2 has been shown to be important for SARS-CoV-1 frameshifting ^37^. In the SL2 mutant, frameshifting was completely abolished and we detected no interaction with ZAP-S **(Fig. 4B, Supplementary Fig. 3B)**. It was previously suggested that ZAP preferentially binds to CG dinucleotides to discriminate between host and viral RNAs ^41, 48^. To test whether ZAP-S binding to SARS-frameshift motif was due to the presence of CG motifs, we generated a control RNA with same nucleotide compositions as the wild-type but lacking CG-dinucleotides. In the CG-depleted control RNA, the predicted pseudoknot fold would not form. ZAP-S interaction was also abolished in the CG-depleted control RNA **(Fig. 4C and Supplementary Fig. 3C)**. In addition, we tested the binding of two control proteins, IGF2BP3 and SUMO, to the SARS-CoV-2 frameshift motif. Compared to ZAP-S, IGF2BP3 as a known RBP showed almost 7X lower affinity to the RNA (K_D_= 806 ± 112 nM). We observed no interaction of the SUMO protein with the frameshift element, confirming the specificity of the RNA-binding assay **(Supplementary Fig. 3D)**. Taken together, these data confirm that ZAP-S preferably binds to elements within the frameshift site and support that those direct interactions with the RNA are crucial for translational regulation.

**Fig. 4.**
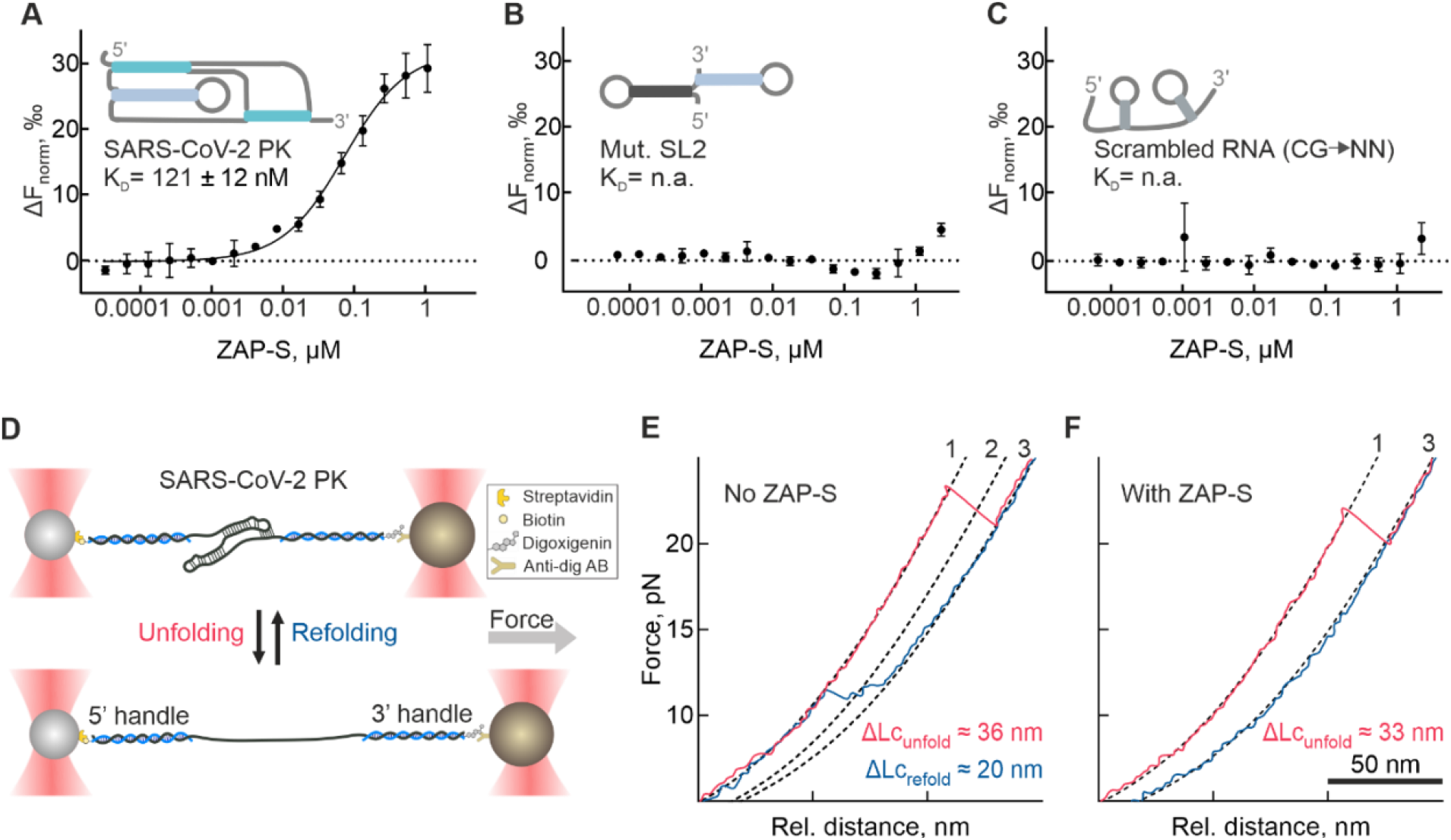
*In vitro* characterization of ZAP-S interaction with SARS-CoV-2 –1 PRF RNA. **(A-C)** Microscale thermophoresis assay to monitor ZAP-S binding to **(A)** SARS-CoV-2 pseudoknot, **(B)** SL2 mutant, **(C)** CG-depleted scrambled (random structure). Unlabeled ZAP-S (40 pM to 2 μM) was titrated against 3’ pCp-Cy5 labeled RNA (5 nM) and thermophoresis was recorded at 25°C with 5% LED intensity and medium MST power. Change in fluorescence (ΔFnorm) was measured at MST on-time of 5 s. Data were analyzed and K_D_ was determined using standard functions in the MO. Affinity Analysis software. Data represent mean ± SD of three measurements (n = 3). (**D)** Schematic illustrating optical tweezers experiments. RNA was hybridized to single-stranded DNA handles flanking the SARS-CoV-2 frameshift site, and conjugated to functionalized beads. A focused laser beam was used to exert pulling force from one end of the molecule. The force was gradually increased until the RNA is fully unfolded (bottom). **(E)** Example unfolding (pink) and refolding (blue) traces of a SARS-CoV-2 RNA sample. **(F)** Example unfolding (pink) and refolding (blue) traces of the SARS-CoV-2 RNA in presence of 400 nM ZAP-S. The numbered and dashed lines represent WLC-based fits (from left to right) of (1) folded, (2) intermediate, and (3) unfolded states as described in Methods. See also **Supplementary Fig. 3 and S4, Supplementary Table 5**.

### ZAP-S prevents the refolding of the stimulatory RNA

We next tested whether the interaction between ZAP-S and the –1PRF RNA alters the RNA structure and mechanical stability. In order to decipher the effect of ZAP-S binding on the folding and unfolding behavior of the frameshifting RNA element, we employed the single-molecule optical-tweezers assay. To this end, a 68 nt long RNA fragment containing the wild type SARS-CoV-2 putative pseudoknot was hybridized to DNA handles. We used the force-ramp method to probe the forces required for unfolding and refolding of the RNA. Briefly, the frameshift RNA was gradually stretched at a constant rate, and then the applied force was released while recording the molecular end-to-end extension distances. This allows the RNA molecule to transition between folded and unfolded states, and sudden changes in measured force-distance trajectories indicate transitions between various RNA conformers **(Fig. 4D)**. With the SARS-CoV-2 putative pseudoknot, in the absence of ZAP-S, we observed two major populations. The first one shows a single long step around 20 pN **(Fig. 4E, Supplementary Table 5)**. At decreasing forces, the RNA refolded in two steps, both at about 11 pN **(Fig. 4E, Supplementary Fig. 4A, Supplementary Table 5)**. Such a hysteresis during refolding is commonly reported with pseudoknots and other highly structured RNAs. Moreover, the contour length change obtained by fitting the data with worm-like chain models was 33.5 ± 3.3 nm **(Supplementary Table 5)**, consistent with the expected value for the full length of the pseudoknot reported previously ^49,50^. We also noted the presence of alternative unfolding trajectories, in line with the alternative RNA conformation, which was recently proposed by the Woodside group for SARS-CoV-2 ^51^. These trajectories were marked with two consecutive unfolding steps occurring at lower forces around 14 pN with the contour length change of 15.9 ± 2.2 nm **(Fig. 4F, Supplementary Fig. 4A, Supplementary Table 5)**. Regardless of the unfolding trajectory, when the force was released, the RNA refolded back in two steps at around 11 pN. Our results were also in line with previous reports on the SARS-CoV-1 frameshift element ^52^. We then carried out the same analysis in the presence of the *trans*-acting protein ZAP-S. To our surprise, in the presence of ZAP-S, the unfolding trajectories of the RNA remained almost unaffected, arguing against an RNA stabilizing effect of the interaction (**Fig. 4F, Supplementary Fig. 4B-D, Supplementary Table 5**). On the other hand, strikingly refolding of the RNA into its native fold was impaired with less or no detectable transitions into the folded state **(Fig. 4F, Supplementary Fig. 4C, D)**. We thus concluded that ZAP-S interaction impairs the refolding of the SARS-CoV-2 frameshift stimulatory RNA, how this works in the presence of ribosomes awaits investigation.

### ZAP-S interacts with translating ribosomes

Recoding events, in general, are dynamic processes that involve the synergistic action of translating ribosomes and mRNA stimulatory elements. As demonstrated by our ensemble and single-molecule analysis, ZAP-S directly interacts with the SARS-CoV-2 frameshift signal and alters folding of the RNA element. However, several *trans*-acting factors including the cardiovirus 2A and SFL were also shown to bind to ribosomes ^15,53^. To explore the possibility of whether ZAP also interacts with ribosomes during translation, we performed ribosome pelleting assay **(Fig. 5A)**. We employed lysates of SARS-CoV-2 infected and uninfected Huh7 cells stably overexpressing ZAP-S. Ribosomes in these lysates were then pelleted through a sucrose cushion, and associated proteins were analyzed via western blotting **(Fig. 5B)**. We were able to demonstrate that ZAP-S was present in the ribosome pellet. Since ZAP-L is constitutively expressed in human tissues, we also detected the presence of endogenous ZAP-L in the ZAP-S overexpressed cells ^33^. To test whether the endogenous ZAP-S is also associated with ribosomes, we similarly processed infected and uninfected naïve Calu-3 cells without any ZAP overexpression **(Supplementary Fig. 5A)**. Likewise, endogenous ZAP-S and ZAP-L could also be easily detected in the ribosome pellet. Additionally, we also detected viral nsp1 protein in the pellet, which is known to associate with the small ribosomal subunit during SARS-CoV-2 infection **(Supplementary Fig. 5A)** ^54^. To test whether ZAP-S is associated with actively translating ribosomes, we also performed polysome profiling using HEK293 cells overexpressing ZAP-S. ZAP-S was detected in most of the ribosomal fractions including polysomes **(Fig. 5C)**. In contrast, the endogenous longer isoform, ZAP-L was identified only in the free RNA fractions. We observed similar polysome profiles with cells overexpressing the SFL, which was bound to the free RNA, the 40S and 60S subunits, the 80S ribosomes, and polysomes **(Supplementary Fig. 5B)** ^15^. Collectively, these results indicate that ZAP-S can be found associated with ribosomes, including the actively translating polysomes. Therefore, we posited the involvement of ZAP in fine-tuning translation. It remains to be investigated, whether this is a physiological phenomenon occurring during different viral infections.

**Fig. 5.**
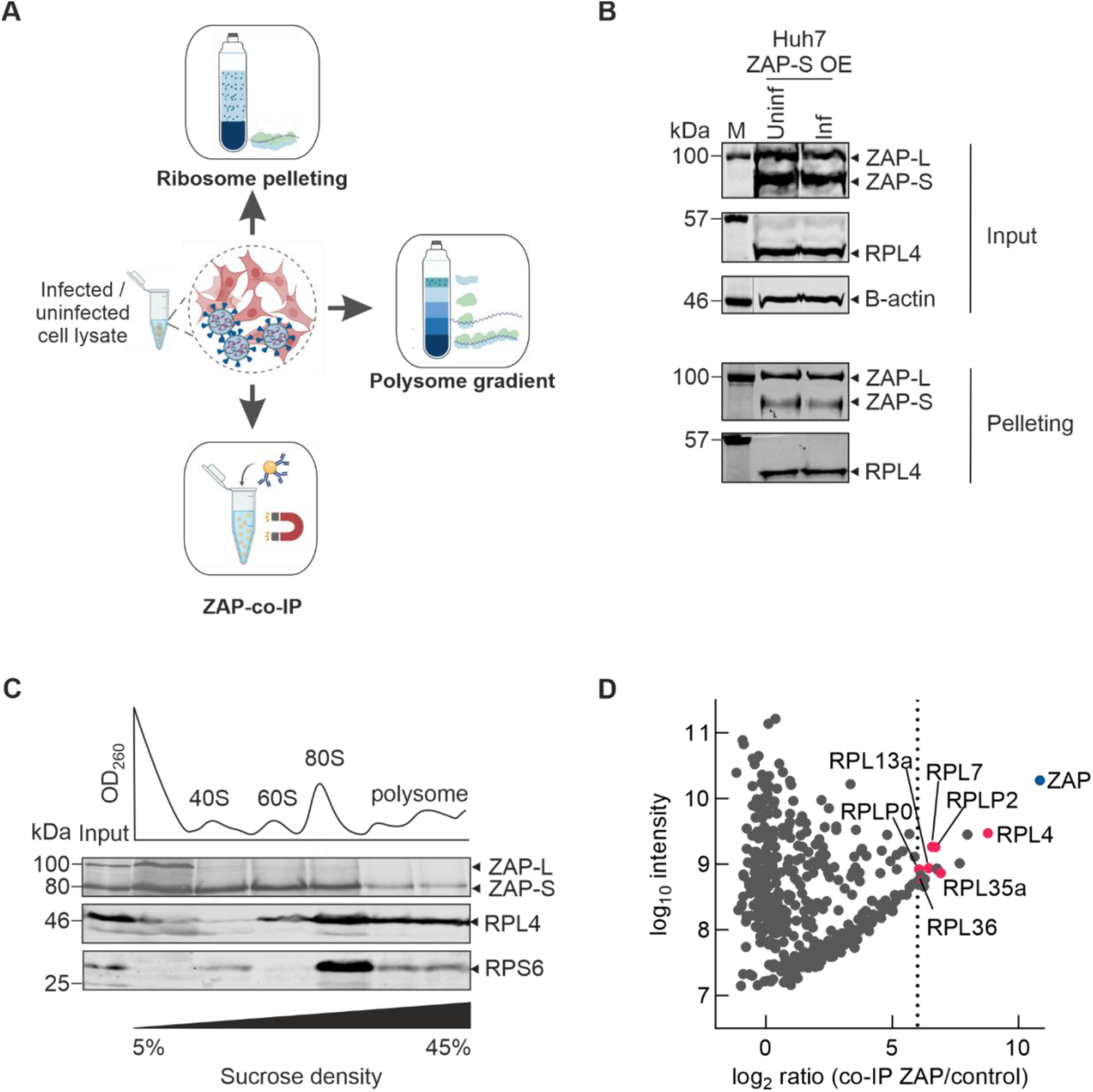
Interactions of ZAP-S with the translation machinery. **(A)** Schematic of the experimental workflow to analyze ribosome association. **(B)** Ribosome pelleting of infected cells. Uninfected and SARS-CoV-2 infected HuH-7 cells stably expressing ZAP-S were lysed and loaded onto sucrose cushions. Levels of ribosomal proteins, ZAP, and B-actin in the pellets were analyzed by western blotting using anti-RPL4, anti-RPS6, anti-ZC3HAV1 (ZAP) and anti-beta actin antibodies. **(C)** Polysome profiling analysis of ZAP. HEK293 cells both naïve and transiently expressing ZAP-S were lysed, subjected to 5-45% sucrose gradient ultracentrifugation, and subsequently fractionated. Levels of ribosomal proteins, as well as ZAP in each fraction, were analyzed by western blotting using anti-RPL4, anti-RPS6, and anti-ZC3HAV1 (ZAP) antibodies. **(D)** Ratio vs. intensity plot of a co-immunoprecipitation of ZAP from SARS-CoV-2-infected Calu-3 lysate. Highlighted proteins were enriched by at least 6-fold in comparison to control rabbit IgG. A full list of hits can be found in **Supplementary Table 3**.

We were surprised to observe strong association of ZAP with RNA and ribosomes, which may provide some clues on how the factor can regulate translation of viral RNAs. To detail physiologically relevant cellular interaction partners of ZAP during infections, we performed co-immunoprecipitation (co-IP) in SARS-CoV-2 infected Calu-3 cells. Captured proteins following affinity purification were detected by LC-MS/MS analysis **(Supplementary Fig. 5C, D, Supplementary Table 2)**. As a control to differentiate between RNA-vs. protein-mediated interactions, we also treated the lysates with Benzonase endonuclease, which degrades nucleic acids. Upon analysis, a total of 124 proteins were detected with at least 4-fold enrichment. The prominence of translation-relevant proteins in untreated and nuclease treated samples was striking. 41% of the enriched proteins were reported to be involved in translation, 48% in protein localization to the ER, 44% in translation initiation, and 41% in ribosome biogenesis **(Supplementary Fig. 5C)**. Previously it was reported that ZAP interacts with proteins including TRIM25 ^39^, eIF4A ^55^ and KHNYN ^40^. In the present co-IP analysis, we were able to detect TRIM21 – a close relative of TRIM25, which supports the interplay between TRIM proteins and ZAP ^56^. We also identified eIF4H as a prominent hit in our analysis, which is the cofactor of the DEAD-box RNA helicase eIF4A ^57^. Furthermore, we detected the RNA-binding protein LARP1 as a ZAP-interaction partner, which may point to a novel route for regulation of translation initiation by ZAP ^58^. As for ribosomal proteins, we found large subunit proteins RPLP0, RPLP2, RPL10A, RPL12, RPS12 and RPS16 among the most enriched interaction partners **(Fig. 5D)**. Intriguingly, in contrast to SFL, which stimulated premature termination, we were not able to detect any translation release factors, pointing to a crucial difference in their mechanisms of action. Taken together, these results indicate that ZAP does not only associate with RNA elements but also with ribosomal proteins through protein-protein mediated interactions. This expands the role of ZAP as a SARS-CoV frameshifting modulator, proposing novel functions beyond its previously suggested roles in inhibition of translation initiation ^55^. Currently, we are further dissecting these interactions using structural analysis tools.

## DISCUSSION

Programmed ribosomal frameshifting (–1PRF) is indispensable for the expression of RNA-dependent RNA polymerase (RdRP) of coronaviruses. In this study, we explored for the first time whether *trans*-factors modulate SARS-CoV-2 –1PRF in infected versus uninfected cells. We discovered that the interferon-induced antiviral protein ZAP-S can strongly impair SARS-CoV-2 frameshifting.

A large number of the interactors found in our screen are well-known RNA-binding proteins and/or regulators of translation. Most of the interactors we identified using the minimal frameshift element as the bait also appeared in genome-wide studies of SARS-CoV-2 RNA binding proteins ^3,30–32^. Despite technical differences and different systems employed among these studies, ZAP-S was surprisingly one of the prominent common hits. We observed that many of the binders were not changing the frameshift levels *in vitro*, which is in line with the notion that mRNA association alone is not sufficient to modulate frameshifting levels. In addition, the mechanism by which ZAP acts on the frameshifting seems to be more complex than anticipated for other *cis-* and *trans*-acting regulators of frameshifting. It is well established that ZAP can exert its antiviral activity by binding to CG-rich RNAs and thereby targeting them for degradation. Yet, ZAP does not bind to all CG-rich sequences and location of these sequences were shown to be crucial for ZAP interaction ^36, 37^. We speculate that, in addition to recognizing CG dinucleotides within the SARS-CoV-2 RNA frameshift site, ZAP-S also recognizes a particular fold within the putative pseudoknot. This way the factor modulates RNA folding pathways and thus plasticity, which is a common feature of RNA-based regulation of biological processes ^59,60^. In fact, structural plasticity has been reported as a hallmark of –1PRF stimulatory RNAs ^47^. Our single-molecule pulling experiments indicate that ZAP-S binds to the SARS-CoV-2 –1PRF RNA directly and interferes with the refolding of the stimulatory RNA structure *in vitro*. As a result, unlike other viral and cellular factors, such as the cardiovirus 2A or poly(C) binding protein, which were shown to directly associate with RNA motifs to induce frameshifting, here association ZAP-S, a host-encoded antiviral protein with the RNA element significantly reduces frameshifting ^12, 14^.

Beyond solely binding the –1PRF stimulatory structure, we were surprised to detect interactions between ZAP-S and the translation machinery. The translation-inhibitory effect of ZAP was previously suggested to be limited to initiation through sequestration of eIF4A. The repression of translation by ZAP was also reported to be stimulated by its co-factor TRIM25 ^39^. In our *in vitro* assays, ZAP-S alone was sufficient to modulate translational frameshifting without additional co-factors. Interaction with ribosomes was observed also with other trans-factors such as the SFL protein ^15^. However, despite the similarities, our data suggests a different mechanism of action for ZAP-S. SFL was shown to interact with stalled ribosomes and recruit release factors to terminate translation, and it acts not only on the HIV-1 but also other frameshift sites including the cellular frameshift gene *PEG10* ^15^. Here, we were not able to detect any release factors by ZAP co-IP analysis, and effect of ZAP-S was specific to SARS-CoV-2 and the closely related SARS-CoV-1. Overall, these findings establish ZAP-S as the first cellular factor, which has a direct role in modulating SARS-coronavirus frameshifting. In accordance with previously published results, we demonstrate that overexpression of ZAP-S reduces the replication of SARS-CoV-2 by 90% ^44, 20^. Further studies are required to deconvolute the multivalent effects of ZAP-S on immunity, viral replication and RNA-related processes such as translation and RNA stability ^33, 38 – 42^.

Ultimately, based on our findings, we propose the following model for the inhibition of –1PRF by ZAP-S **(Fig. 6)**. Accordingly, ZAP-S binding to the frameshift RNA can alter the stimulatory RNA structure and reduce the chance of elongating ribosomes to encounter the stimulatory pseudoknot. Without a stimulatory structure, the elongation pause during next round of translation would be too short for codon-anti-codon interactions to be established in the –1 frame. Thus, ZAP-S would likely allow translation to proceed without any roadblock effect and terminate at the 0-frame UAA stop codon found immediately downstream of the slippery sequence. How exactly the binding of ZAP-S and other riboregulators to ribosomes would impact kinetics of translation and frameshifting awaits further investigation.

**Fig. 6.**
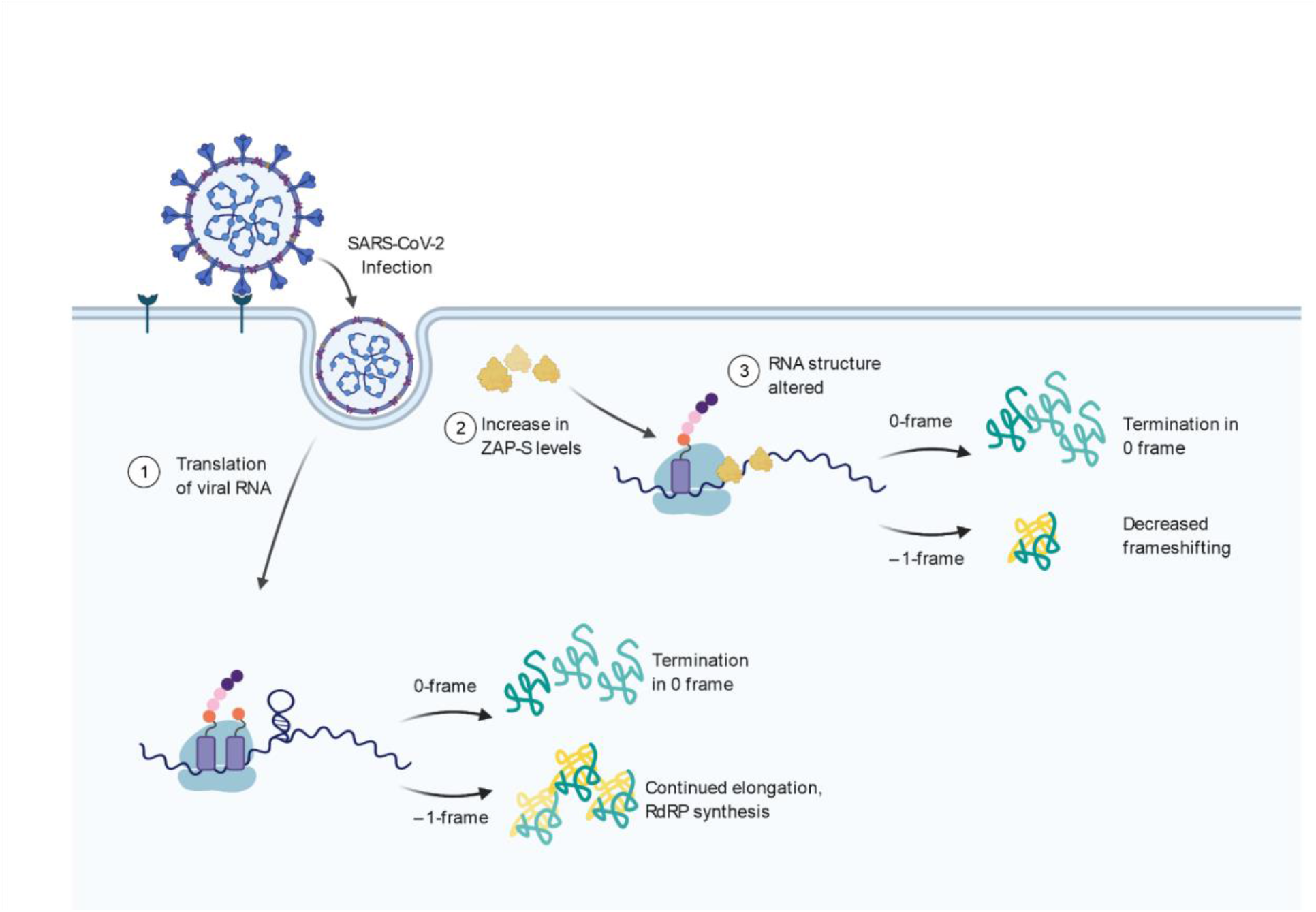
Model for ZAP-S mediated inhibition of SARS-CoV-2 frameshifting. **(1)** Upon infection, the viral RNA is translated by the cellular machinery, where 40% of translation events yield the 1a/1b polyprotein through –1PRF. **(2)** Infection also leads to the induction of antiviral factors including ZAP-S. **(3)** ZAP-S binding to the frameshift RNA alters the RNA structure and reduces the chance of elongating ribosomes to encounter the stimulatory structure. In absence of this structure, the elongation pause would be too short for codon-anti-codon interactions to be established in the –1 frame. Thus, ZAP-S would allow translation to proceed without any roadblock effect and terminate at the canonical 0-frame UAA stop codon found just downstream of the slippery sequence. This would lead to a decrease in the amounts of the 1a/1b polypeptides and thus hamper the production of the RdRP from the –1 frame.

Given the plethora of mechanisms presented by which *trans*-regulators of PRF can act, it is conceivable that viral and host encoded trans-factors follow a multitude of routes to impact frameshift paradigms. Taken together, this study highlights a novel role for ZAP-S in SARS-CoV-2 antiviral response and establishes ZAP-S as a *bona fide* regulator of SARS-CoV-2 translation. This work also provokes the idea of using ZAP-S as a potential SARS-CoV-2 intervention strategy, by employing ZAP-S mRNA as a multifunctional antiviral and immune-modulatory drug.

## MATERIALS AND METHODS

### RNA affinity pulldown mass spectrometry

RNA antisense purification was performed according to a protocol based on ^17^. Briefly, 6*10^7 HEK293 cells per condition were lysed in a buffer containing 20 mM Tris/HCl pH 7.5, 100 mM KCl, 5 mM MgCl2, 1 mM DTT, 0.5 % Igepal CA630 (Sigma-Aldrich), 1× cOmplete^™^ Protease Inhibitor Cocktail (Roche), 40 U/ml RNase inhibitor (Molox). The cleared lysate was incubated with *in vitro* transcribed RNA corresponding to the SARS-CoV-2 –1PRF site, which was immobilized on streptavidin hydrophilic magnetic beads (NEB) by biotin-streptavidin interaction. After three washes with binding buffer (50 mM HEPES/KOH pH 7.5, 100 mM NaCl, 10 mM MgCl2) and two washes with wash buffer (50 mM HEPES/KOH pH 7.5, 250 mM NaCl, 10 mM MgCl2), bound proteins were eluted by boiling the sample in 1× NuPAGE LDS sample buffer (Thermo Fisher Scientific) supplemented with 40 mM DTT. For infected as well as uninfected Calu-3 cells the procedure was performed similarly. In order to inactivate the virus, the lysis buffer contained Triton-X100 and inactivation was confirmed by plaque assays.

For LC-MS/MS, the eluted proteins were alkylated using iodoacetamide followed by acetone precipitation. In solution digests were performed in 100 mM ammonium bicarbonate and 6 M urea using Lys-C and after reducing the urea concentration to 4 M with trypsin. Peptides were desalted using C18 stage tips and lyophilized. LC-MS/MS was performed at the RVZ Proteomics Facility (Würzburg) and analyzed as described previously ^61^. Gene ontology (GO) term analysis was performed with Panther ^62^. The list of all identified proteins is given in **Supplementary Table 1**.

### Co-immunoprecipitation

Endogenous interaction partners of ZAP were identified by co-immunoprecipitation followed by mass spectrometry as published previously ^63^. Briefly, uninfected and SARS-CoV-2-infected Calu-3 cells were lysed in lysis buffer (10 mM Tris/HCl pH 7.4, 150 mM NaCl, 1% Igepal CA630 (Sigma-Aldrich) and 1x cOmplete^™^ Protease Inhibitor Cocktail (Roche)). The lysis buffer was supplemented either with 40 U/ml RNase inhibitor (Molox) or 50 U/ml Benzonase (Roche) to differentiate between RNA- and protein-dependent interactions. 1 mg of cell lysate was cleared with protein A magnetic beads to remove non-specific interactions (S1425S, NEB) and incubated overnight with anti-ZAP antibody (Proteintech 16820-1-AP) or anti-IgG from rabbit as a control (Cell Signaling, a gift from Dr. Mathias Munschauer, HIRI-HZI). Antibodies were captured with protein A magnetic beads, washed with lysis buffer, and eluted by boiling in 1X NuPAGE^™^ LDS Sample Buffer (Thermo Fisher Scientific) supplemented with 40 mM DTT. LC-MS/MS proceeded as described above. A list of all identified proteins can be found in **Supplementary Table 2**.

### Plasmid construction

To generate dual-fluorescence reporter constructs frameshift sites of SARS-CoV, SARS-CoV-2, MERS-CoV, BtCoV 273, HIV-1, JEV, PEG10, WNV were placed between the coding sequence of EGFP and mCherry (parental construct was a gift from Andrea Musacchio (Addgene plasmid # 87803 ^64^) by site-directed mutagenesis in a way that EGFP would be produced in 0-frame and mCherry in –1-frame. EGFP and mCherry were separated by StopGo ^65^ signals as well as an alpha-helical linker ^66^. A construct with no PRF insert and mCherry in-frame with EGFP served as a 100% translation control and was used to normalize EGFP and mCherry intensities.

To generate screening vectors, protein-coding sequences of DDX17 (NM_001098504.2), DDX36 (NM_020865.3), ELAVL1 (NM_001419.3), GNL2 (NM_013285.3), HNRNPF (NM_001098204.2), HNRNPH1 (NM_001364255.2), HNRNPH2 (NM_001032393.2), IGF2BP1 (NM_006546.4), MATR3 iso 2 (NM_018834.6), MMTAG2 (NM_024319.4), NAF1 (NM_138386.3), NHP2 (NM_017838.4), POP1 (NM_001145860.2), RAP11B (NM_004218.4), RSL1D1 (NM_015659.3), SFL (NM_018381.4), SURF6 (NM_001278942.2), TFRC (NM_003234.4), ZAP (NM_024625.4) and ZNF346 (NM_012279.4) were placed in frame with the coding sequence for ECFP in pFlp-Bac-to-Mam (gift from Dr. Joop van den Heuvel, HZI, Braunschweig, Germany ^67^) via Gibson Assembly ^68^.

Golden Gate compatible vectors for heterologous overexpression in *E. coli, in vitro* translation in RRL, and lentivirus production, were generated by Golden Gate or Gibson Assembly. A dropout cassette was included to facilitate the screening of positive colonies. Protein-coding sequences were introduced by Golden Gate Assembly using AarI cut sites ^69^. pET-SUMO-GFP (gift from Prof. Utz Fischer, Julius-Maximilians-University, Würzburg, Germany) was used as the parental vectors for protein overexpression in *E. coli*. The lentivirus plasmid was a gift from Prof. Chase Beisel (HIRI-HZI, Würzburg, Germany). An ALFA-tag was included to facilitate the detection of the expressed protein ^70^. The frameshift reporter vector for the *in vitro* translation contained ß-globin 5’ and 3’ UTRs as well as a 30 nt long poly(A) tail. The insert was derived from nucleotides 12686–14190 of SARS-CoV-2 (NC_045512.2); a 3×FLAG-tag was introduced at the N-terminus to facilitate detection. To generate 0% and 100% –1PRF controls, the –1PRF site was mutated by disrupting the pseudoknot structure as well as the slippery sequence.

Optical tweezers constructs were based on the wild type SARS-CoV-2 frameshifting site (nucleotides 13475-13541) cloned into the plasmid pMZ_lambda_OT, which encodes for the optical tweezer handle sequences (2Kb each) flanking the RNA structure (130 nt). Constructs were generated using Gibson Assembly. Sequences of all plasmids and oligos used in this study are given in **Supplementary Table 4**.

### Cell culture, transfections, generation of polyclonal stable cell lines

HEK293 cells (gift from Prof. Jörg Vogel, HIRI-HZI) and Huh7 cells (gift from Dr. Mathias Munschauer, HIRI-HZI), were maintained in DMEM (Gibco) supplemented with 10% FBS (Gibco) and 100 U/ml streptomycin and 100 mg/ml penicillin. Calu-3 cells (ATCC HTB-55) were cultured in MEM (Sigma) supplemented with 10% FBS. Cell lines were kept at 37 °C with 5% CO2. Transfections were performed using PEI (Polysciences) according to manufacturer’s instructions. For co-transfections, plasmids were mixed at a 1:1 ratio.

VSV-G envelope pseudo-typed lentivirus for the generation of stable cell lines was produced by co-transfection of each transfer plasmid with pCMVdR 8.91 ^71^ and pCMV-VSV-G (gift from Prof. Weinberg, Addgene plasmid # 8454 ^72^). 72 h post-transfection, the supernatant was cleared by centrifugation and filtration. The supernatant was used to transduce naïve Huh7 cells in the presence of 10 μg/ml polybrene (Merck Millipore). After 72 h, the cells were selected with 10 μg/ml blasticidin (Cayman Chemical) for 10 days to generate polyclonal cell lines.

### SARS-CoV-2 infection

For infection with SARS-CoV-2, we used the strain hCoV-19/Croatia/ZG-297-20/2020, a kind gift of Prof. Alemka Markotic (University Hospital for Infectious Diseases, Zagreb, Croatia). The virus was raised for two passages on Caco-2 cells (HZI Braunschweig). Calu-3 cells (ATCC HTB-55) were infected with 2000 PFU/ml corresponding to an MOI of 0.03 at 24h post-infection, cells were collected and lysed for proteomic and ribosome-interaction experiments. To study the effect of ZAP-S on SARS-CoV-2 infection, Huh-7 cells were employed. One hour before infection, Huh-7 cells both naïve or ZAP-S-overexpressing cells were either pre-stimulated with IFN-β (500 U/ml), IFN-γ (500 U/ml), IFN-λ1 (5 ng/ml), or left untreated. Cells were infected with 200000 pfu/ml, corresponding to an MOI of 0.03 at 24h post-infection, cell culture supernatants were collected and titrated by plaque assay on Vero E6 cells (ATCC CRL-1586). Briefly, confluent Vero E6 cells in 96-well plates were inoculated with dilutions of the virus-containing supernatants for one hour at 37 °C, the inoculum was removed and cells were overlaid with MEM containing 1.75% methyl-cellulose. At three days post-infection, whole wells of the plates were imaged using an IncuCyte S3 (Sartorius) at 4x magnification, and plaques were counted visually.

### Flow cytometry

HEK293 cells were transiently transfected with either the control construct or the –1PRF construct encoding for the dual-fluorescence EGFP-mCherry translation reporter as outlined in **Fig. 2A**. Cells were harvested at 24 h post-transfection and fixed with 0.4% formaldehyde in PBS. After washing with PBS, flow cytometry was performed on a FACSAria III (BD Biosciences) or a NovoCyte Quanteon (ACEA) instrument. Flow cytometry data were analyzed with FlowJo software (BD Biosciences). ECFP-positive cells were analyzed for the ratio between mCherry and EGFP **(Supplementary Fig. 2F)**. FE was calculated according to the following formula:

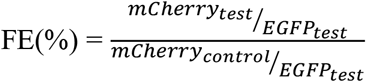

where mCherry represents the mean mCherry intensity, EGFP the mean EGFP intensity, test represent the tested sample and control represents the in-frame control where mCherry and EGFP are produced in an equimolar ratio ^73^. Data represent the results of at least three independent experiments.

### Purification of recombinant proteins

Recombinant ZAP-S N-terminally tagged with 6×His-SUMO was purified from *E. coli* Rosetta 2 cells (Merck) by induction with 0.2 mM isopropyl β-d-1-thiogalactopyranoside for 18 h at 18 °C. Cells were collected, resuspended in lysis buffer (50 mM HEPES/KOH pH 7.6, 1 M NaCl, 1 mM DTT, 1 mM PMSF) and lysed in a pressure cell. The lysate was cleared by centrifugation and ZAP-S was captured using Ni-NTA resin (Macherey-Nagel). After elution with 500 mM imidazole, ZAP-S was further purified and the bound nucleic acids removed by size exclusion chromatography (HiLoad^®^ 16/600 Superdex^®^ 200) in 20 mM HEPES/KOH pH 7.6, 1 M KCl, 1 mM DTT, 20% glycerol. Protein identity was verified by SDS-PAGE as well as western blotting (**Supplementary Fig. 2D)**. Purified ZAP-S was rapidly frozen and stored in aliquots at −80 °C.

His-SUMO-IGF2BP3 as well as His-SUMO were kind gifts from Dr. Andreas Schlundt (Goethe University, Frankfurt, Germany)

### Western blots

Protein samples were denatured at 95 °C and resolved by 12 % SDS-PAGE at 30 mA for 2 h. After transfer using Trans-Blot (Bio-Rad), nitrocellulose membranes were developed using the following primary antibodies: anti-His-tag (ab18184), anti-DDDDK (ab49763), anti-ALFA (FluoTag^®^-X2 anti-ALFA AlexaFluor 647), anti-ZC3HAV1 (Proteintech 16820-1-AP). The following secondary antibodies were used: IRDye^®^ 800CW Goat anti-rabbit and IRDye^®^ 680RD Donkey anti-Mouse (both LI-COR). Bands were visualized using an Odyssey Clx infrared imager system (LI-COR) or a Typhoon7000 (GE Healthcare).

### *In vitro* translation assays

mRNAs were *in vitro* transcribed using T7 polymerase purified in-house using linearized plasmid DNA as the template. These mRNAs were capped (Vaccinia Capping System, NEB) and translated using the nuclease-treated rabbit reticulocyte lysate (RRL; Promega). Typical reactions were comprised of 75% v/v RRL, 20 μM amino acids, and were programmed with ~50 μg/ml template mRNA. ZAP-S was titrated in the range of 0-3 μM. Reactions were incubated for 1 h at 30 °C. Samples were mixed with 3X volumes of 1X NuPAGE^™^ LDS Sample Buffer (Invitrogen), boiled for 3 min, and resolved on a NuPAGE^™^ 4 to 12% Bis-Tris polyacrylamide gel (Invitrogen). The products were detected using western blot (method as described above). The nitrocellulose membranes were developed using anti-DDDDK primary (Abcam ab49763) and IRDye^®^ 680RD donkey anti-mouse secondary antibody (LI-COR). Bands were visualized using an Odyssey Clx infrared imager system (LI-COR). Bands corresponding to the –1 or 0-frame products, 58 kDa and 33 kDa respectively, on western blots of *in vitro* translations were quantified densitometrically using ImageJ software ^74^. FE was calculated as previously described, by the formula intensity (–1-frame)/ (intensity (–1-frame) + intensity (0-frame)) ^11^. The change in FE was calculated as a ratio of FE of each condition to the FE of no-protein control in each measurement. Experiments were repeated at least 3 independent times.

### Microscale thermophoresis

Short frameshifting RNA constructs were *in vitro* transcribed using T7 polymerase as described above. RNAs were labeled at the 3’ end using pCp-Cy5 (Cytidine-5’-phosphate-3’-(6-aminohexyl) phosphate) (Jena Biosciences). For each binding experiment, RNA was diluted to 10 nM in Buffer A (50 mM Tris-HCl pH 7.6, 250 mM KCl, 5 mM MgCl2, 1 mM DTT, 5% glycerol supplemented with 0.05% Tween 20 and 0.2 mg/ml *E. coli* tRNA). A series of 16 tubes with ZAP-S dilutions were prepared in Buffer A on ice, producing ZAP-S ligand concentrations ranging from 40 pM to 2 μM. For measurements, each ligand dilution was mixed with one volume of labeled RNA, which led to a final concentration of 5.0 nM labeled RNA. The reaction was mixed by pipetting, incubated for 10 min at room temperature, followed by centrifugation at 10,000 × g for 5 min. Capillary forces were used to load the samples into Monolith NT.115 Premium Capillaries (NanoTemper Technologies). Measurements were performed using a Monolith Pico instrument (NanoTemper Technologies) at an ambient temperature of 25 °C. Instrument parameters were adjusted to 5% LED power, medium MST power, and MST on-time of 5 seconds. An initial fluorescence scan was performed across the capillaries to determine the sample quality and afterward, 16 subsequent thermophoresis measurements were performed. Data of three independently pipetted measurements were analyzed for the ΔFnorm values, and binding affinities were determined by the MO. Affinity Analysis software (NanoTemper Technologies). Graphs were plotted using GraphPad Prism 8.4.3 software.

### Microscopy

HEK293 cells were cultured on glass slides and transfected as described above. The cells were fixed with 4% paraformaldehyde in 1x PBS for 15 min at room temperature. After washing with 1x PBS, cells were mounted in ProLong Antifade Diamond without DAPI (Invitrogen). Microscopy was performed using a Thunder Imaging System (Leica) using 40% LED power and the 40x objective. EGFP was excited at 460-500 nm and detected at 512-542 nm. mCherry was excited at 540-580 nm and detected at 592-668 nm. The images were processed with the LasX software (Leica).

### Polysome profiling analysis

A plasmid expressing ZAP-S N-terminally tagged with a His-tag was transfected into HEK293 cells using PEI, as described above. To check endogenous ZAP-S expression, HEK cells were transfected with a plasmid containing the same backbone and His-tag. At 24 h post-transfection, cycloheximide (VWR) was added to the medium at a final concentration of 100 μg/ml to stop translation. Approximately 107 HEK cells were lysed with 500 μl lysis buffer (20 mM Tris-HCl pH 7.4, 150 mM NaCl, 5 mM MgCl2, 1 mM DTT, 100 μg/ml Cycloheximide, 1% Triton X), and the lysate was clarified by centrifugation at 17,0000 × g for 10 min at 4 °C. Polysome buffer (20 mM Tris-HCl pH 7.4, 150 mM NaCl, 5 mM MgCl2, 1 mM DTT, 100 μg/ml Cycloheximide) was used to prepare all sucrose solutions. Sucrose density gradients (5%–45% w/v) were freshly made in SW41 ultracentrifuge tubes (Beckman) using a Gradient Master (BioComp Instruments) according to manufacturer’s instructions. The lysate was then applied to a 4%–45% sucrose continuous gradient and centrifuged at 35,000 rpm (Beckmann Coulter Optima XPN) for 3 h, at 4 °C. The absorbance at 254 nm was monitored and recorded and 500 μl fractions were collected using a gradient collector (BioComp instruments). The protein in each fraction was pelleted with trichloroacetic acid, washed with acetone, and subjected to western blotting, as described above.

### Ribosome pelleting assay

SARS-CoV-2 Calu-3 infected lysates were prepared as described above. 300 μl of the lysate was loaded onto a 900 μl 1M sucrose cushion in polysome buffer (described above) in Beckman centrifugation tubes. Ribosomes were pelleted by centrifugation at 75,000 rpm for 2 h, at 4 °C, using a Beckmann MLA-130 rotor (Beckman Coulter Optima MAX-XP). After removing the supernatant, ribosome pellets were resuspended in polysome buffer and was used for western blotting, as described above.

### Optical tweezers constructs

5’ and 3’ DNA handles, and the template for *in vitro* transcription of the SARS-CoV-2 putative pseudoknot RNA were generated by PCR using the pMZ_lambda_OT vector. The 3’ handle was labeled during the PCR using a 5’ digoxigenin-labeled reverse primer. The 5’ handle was labeled with Biotin-16-dUTP at the 3’ end following PCR using T4 DNA polymerase. The RNA was *in vitro* transcribed using T7 RNA polymerase. Next, DNA handles (5’ and 3’) and *in vitro* transcribed RNA were annealed in a mass ratio 1:1:1 (5 μg each) by incubation at 95 °C for 10 min, 62 °C for 2 h, 52 °C for 2 h and slow cooling to 4 °C in annealing buffer (80% formamide, 400 mM NaCl, 40 mM HEPES, pH 7.5, and 1 mM EDTA, pH 8) to yield the optical tweezer suitable construct **(Fig. 4E)**. Following the annealing, samples were concentrated by ethanol precipitation, pellets were resuspended in 40 μl RNase-free water, and 4 μl aliquots were stored at −20 °C until use.

### Optical tweezers data collection and analysis

Optical tweezers measurements were performed using a commercial dual-trap platform coupled with a microfluidics system (C-trap, Lumicks). For the experiments, optical tweezers (OT) constructs were mixed with 4 μl of polystyrene beads coated with antibodies against digoxigenin (AD beads, 0.1% v/v suspension, Ø 1.76 μm, Spherotech), 10 μl of assay buffer (20 mM HEPES, pH 7.6, 300 mM KCl, 5 mM MgCl2, 5 mM DTT and 0.05% Tween) and 1 μl of RNase inhibitor. The mixture was incubated for 20 min at room temperature in a final volume of 19 μl and subsequently diluted by the addition of 0.5 ml assay buffer. Separately, 0.8 μl of streptavidin-coated polystyrene beads (SA beads, 1% v/v suspension, Ø 2 μm, Spherotech) were mixed with 1 ml of assay buffer. The flow cell was washed with the assay buffer, and suspensions of both streptavidin beads and the complex of OT construct with anti-digoxigenin beads were introduced into the flow cell. During the experiment, an anti-digoxigenin (AD) bead and a streptavidin (SA) bead were trapped and brought into proximity to allow the formation of a tether. The beads were moved apart (unfolding) and back together (refolding) at a constant speed (0.05 μm/s) to yield the force-distance (FD) curves. The stiffness was maintained at 0.31 and 0.24 pN/nm for trap 1 (AD bead) and trap 2 (SA bead), respectively. For experiments with ZAP-S protein, recombinantly expressed ZAP-S was diluted to 400 nM in assay buffer and introduced to the flow cell. FD data were recorded at a rate of 78125 Hz.

Raw data files were processed using our custom-written python algorithm called Practical Optical Tweezers Analysis TOol (POTATO, https://github.com/lpekarek/POTATO.git, manuscript in preparation). In brief, raw data were first down sampled by a factor of 20 to speed up subsequent processing, and the noise was filtered using Butterworth filter (0.05 filtering frequency, filter order 2). Numerical time derivation was calculated separately for force and distance data. These derivations were statistically analyzed to identify the folding events and their coordinates. For data fitting, we employed a combination of two worm-like chain models (WLC1 for the fully folded double-stranded parts and WLC2 for the unfolded single-stranded parts) as described previously ^53^. Firstly, the initial contour length of the folded RNA was set to 1231 nm, and the persistence length of the double-stranded part was fitted ^53^. Then, the persistence length of the unfolded RNA was set to 1 nm, and the contour length of the single-stranded part was fitted. Data were statistically analyzed, and the results were plotted using Prism 8.0.2 (GraphPad).

### qRT-PCR

Total RNA was isolated as described previously ^75^, and the reverse transcription using RevertAid (Invitrogen) was primed by oligo(dT). Reactions of quantitative real-time PCR (qRT-PCR) were set up using POWER SYBR green Master-mix (Invitrogen) according to manufacturer’s instructions and analyzed on the CFX96 Touch Real-Time PCR Detection System (Bio-Rad) under the following cycling condition: 50 °C for 2 min, 95 °C for 2 min, followed by 40 cycles of 95 °C for 15 s and 60 °C for 30 s, and ending with a melt profile analysis. The fold change in mRNA expression was determined using the 2-ΔΔCt method relative to the values in uninfected samples, after normalization to the housekeeping gene (geometric mean) GAPDH. Statistical analysis was conducted using an unpaired two-tailed *t*-test with Welch’s correction comparing delta Ct values of the respective RNA in uninfected and infected cells. The results were plotted using Prism 8.0.2 (GraphPad).

### Quantification and statistical analysis

All statistical analyses and software used have been mentioned in the Figure Legends and Materials & Methods. Measurements from the *in vitro* western blot assay and *in vivo* dual fluorescence assay resulted from 3 technical replicates. Measurements from single-molecule experiments resulted from a specified number (n) of traces from a single experiment. For the ensemble MST analysis, all analysis from 3 individual replicates was performed in Nanotemper MO. Affinity software.

## Acknowledgments

We thank Dr. Zeljka Macak-Safranko and Prof. Alemka Markotic (University of Zagreb) for providing the SARS-CoV-2 virus isolate prior to publication. We thank Dr. Andreas Schlundt for kind gifts of IGF2BP3 and SUMO proteins (Goethe University, Frankfurt, Germany). We thank Dr. Joop van den Heuvel (HZI) for his suggestions on ZAP-S purification. We thank Prof. Redmond Smyth, Prof. Jörg Vogel, Prof. Lars Dölken, Prof. Utz Fischer and Prof. Thomas Pietschmann for critical reading of the manuscript. We thank expert technical assistance by Tatyana Koch (HIRI-HZI). We thank Stefan Buck for help with optical tweezer data acquisition. We thank Ayse Barut for cell maintenance for infection studies (HZI). We thank Dr. Andreas Schlosser and Stephanie Lamer from the Rudolf Virchow Center for the LC-MS/MS analysis. Figures were partially generated using BioRender.com.

## Funding

This project is funded fully or in part by the

Helmholtz Association

MWK Niedersachsen through Grant Nr. 14-76103-184 CORONA-2/20.

NC received funding from the European Research Council (ERC) Grant Nr. 948636.

## Author contributions

Conceptualization: MZ, AK, NC

Methodology: MZ, AK, NC, UR, LCS

Investigation: MZ, AK, UR, LP Visualization: MZ, AK, NC Supervision: NC, LCS

Writing - original draft: MZ, AK, NC

Writing-review & editing: MZ, AK, NC

## Data and materials availability

All data are available in the main text, the supplementary materials as well as in Mendeley Data: DOI: 10.17632/c7rbxb86k2.1

---

**Supplementary Fig. 1.**
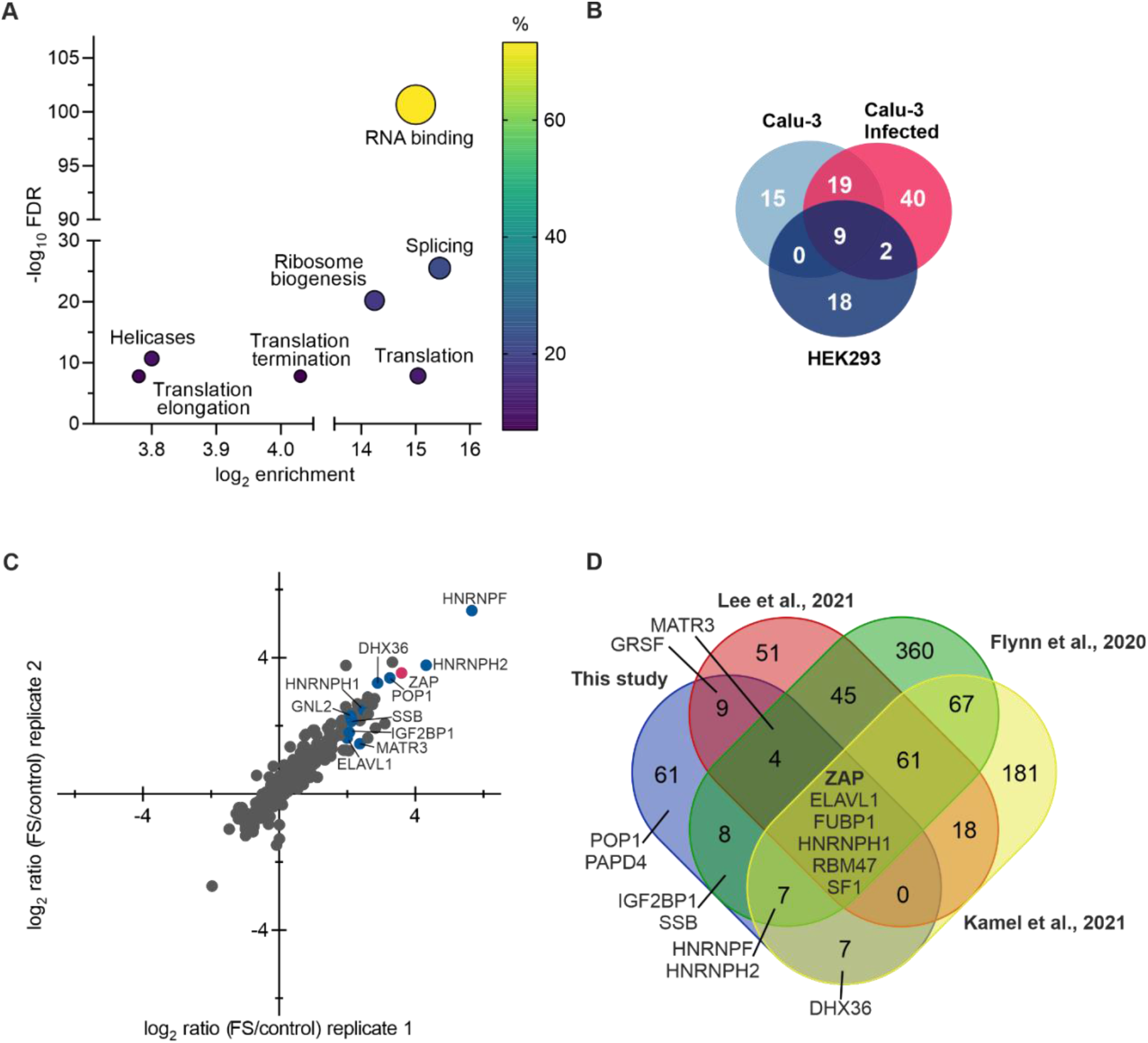
Capture and analysis of frameshift RNA interactors, related to Fig. 1. **(A)** Venn Diagram comparing the hits of the in vitro RNA antisense purification in HEK293, uninfected Calu-3 as well as SARS-CoV-2 infected Calu-3 cells. **(B)** Gene ontology (GO) term analysis of SARS-CoV-2 frameshift RNA interactions. FDR – false discovery rate. **(C)** Scatter plot of log_2_-transformed ratio of RNA-antisense purification in HEK293 cells. **(D)** Venn Diagram comparing the hits of the vitro RNA antisense purification of the SARS-CoV-2 frameshift site from this study with the hits of genome-wide interactome captures from the literature ^3, 20, 21, 22^.

**Supplementary Fig. 2.**
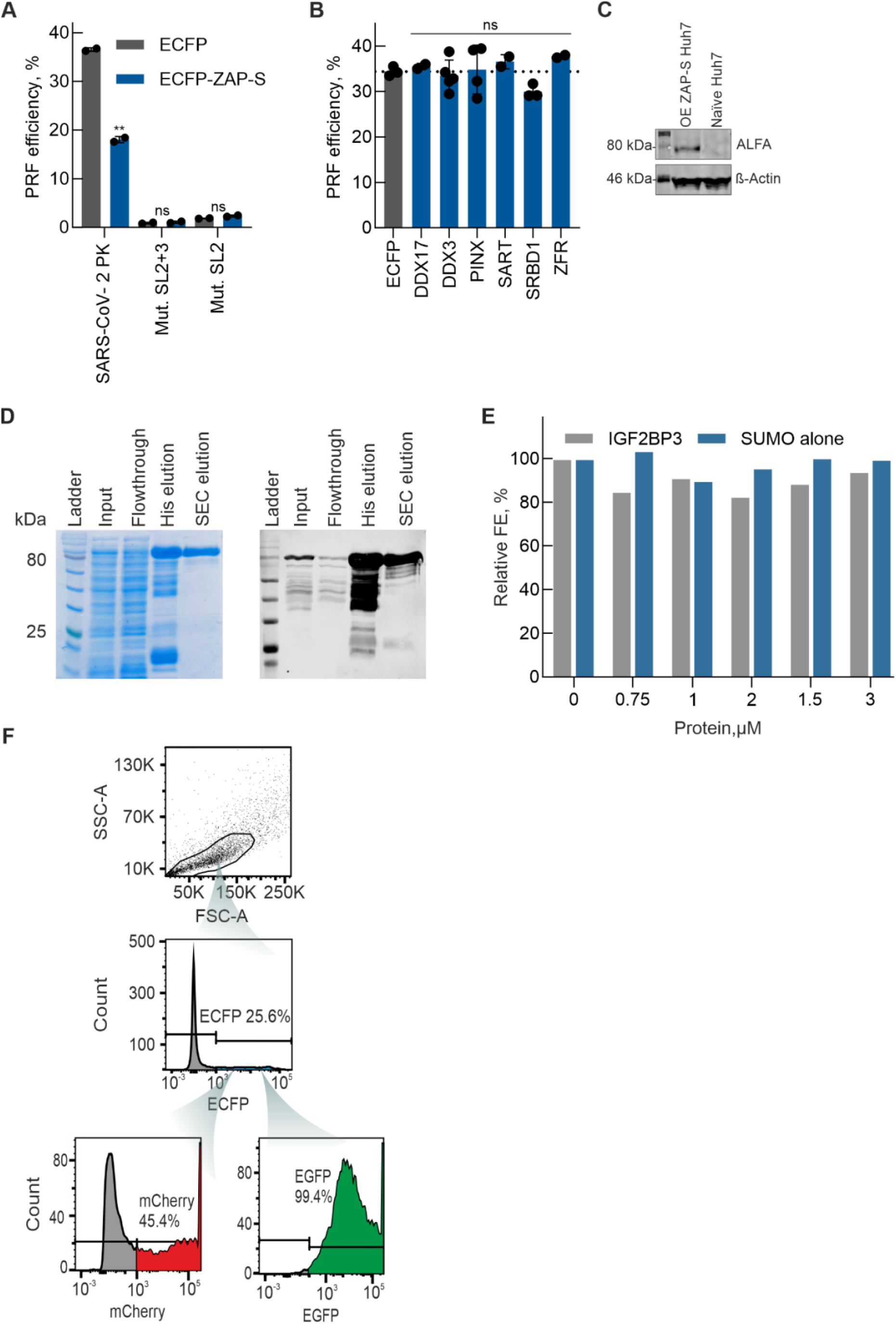
Effect of ZAP-S on FE of various PRF sites, related to Fig. 2 and 3. **(A)** *In vivo* dual-fluorescence of mutants of SARS-CoV-2 –1PRF RNA in HEK293 cells in the presence and absence of ZAP-S. Datapoints represent the mean ± s.d. (n = 2 independent experiments). P values were calculated using an ordinary unpaired one-sided ANOVA comparing values of the ECFP control. * P < 0.01 – ** P < 0.001. See also Table 2. **(B)** *In vivo* dual-fluorescence of SARS-CoV-2 –1PRF RNA in HEK293 cells in the presence and absence of not significantly enriched proteins. Datapoints represent the mean ± s.d. (n = 3 independent experiments). P values were calculated using an ordinary unpaired one-sided ANOVA comparing values of the ECFP control. **(C)** Western Blot of naïve as well as ALFA-tagged ZAP-S-overexpressing Huh7 cells. ALFA-ZAP-S was detected using anti-ALFA antibody, ß-actin serves as a loading control **(D)** Coomassie-stained SDS-PAGE and western blot of heterologous expression of ZAP-S in *E. coli* as well as the purification steps. ZAP-S was detected using an anti-ZC3HAV1 antibody. **(E)** Effect of IGF2BP3 and SUMO on 1a/1b –1 frameshifting *in vitro*. FLAG-tagged SARS-CoV-2 frameshift RNA was translated in RRL in the presence of increasing concentrations of respective protein ranging from 0 to 3 μM. FLAG-tagged peptides generated by ribosomes that do not frameshift (no –1PRF) or that enter the –1 reading frame (−1PRF) were identified via western blotting using anti-DDDDK antibody. Changes in FE observed in the presence of the protein (normalized to 0 μM protein). **(F)** Gating strategy for flow cytometry determining FE in HEK293 cells. Cell populations were determined based on SSC and FSC. ECFP-positive cells were analyzed for the mean intensities of EGFP and mCherry.

**Supplementary Fig. 3.**
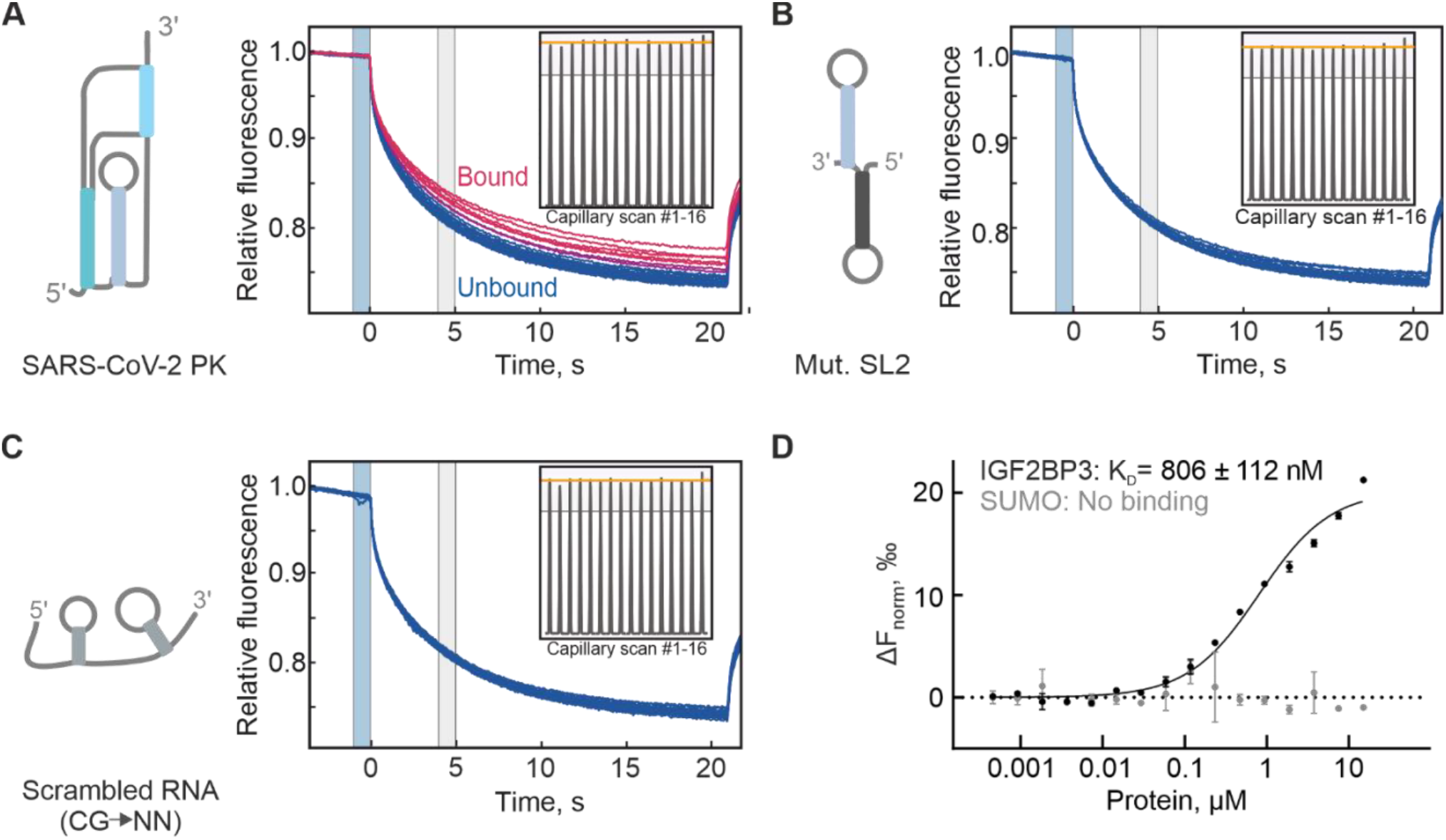
Thermophoresis raw data and interaction of IGF2BP3 and SUMO with SARS-CoV-2 FS RNA, related to Fig. 4. Capillary scans and thermophoretic time-traces of microscale thermophoresis (MST) measurements of binding between ZAP-S and (A) SARS-CoV-2 pseudoknot, (B) SL2 mutant and (C) GC-depleted scrambled (random structure). The grey boxes in the capillary scans mark 20% above and below the average peak fluorescence, the acceptable limit of deviations across the fluorescence scans. Blue and grey boxes in the time-course traces represent the temperature jump and MST-on time (5s), respectively. In all cases, there is no adsorption of the labeled protein to the capillaries. See Fig. 4 for resulting binding curves. (D) Microscale thermophoresis to monitor binding of IGF2BP3 and SUMO to SARS-CoV-2 FS PK. Unlabeled protein (0.4 nM to 15 μM) was titrated against 3’ pCp-Cy5 labeled RNA (5 nM) and thermophoresis was recorded at 25°C with 5% LED intensity and medium MST power. Change in fluorescence (ΔFnorm) was measured at MST on-time of 5s. Data were analyzed and Kd was determined using standard functions in the MO. Affinity Analysis software. Data represent mean +/-SD of two measurements (n=2).

**Supplementary Fig. 4.**
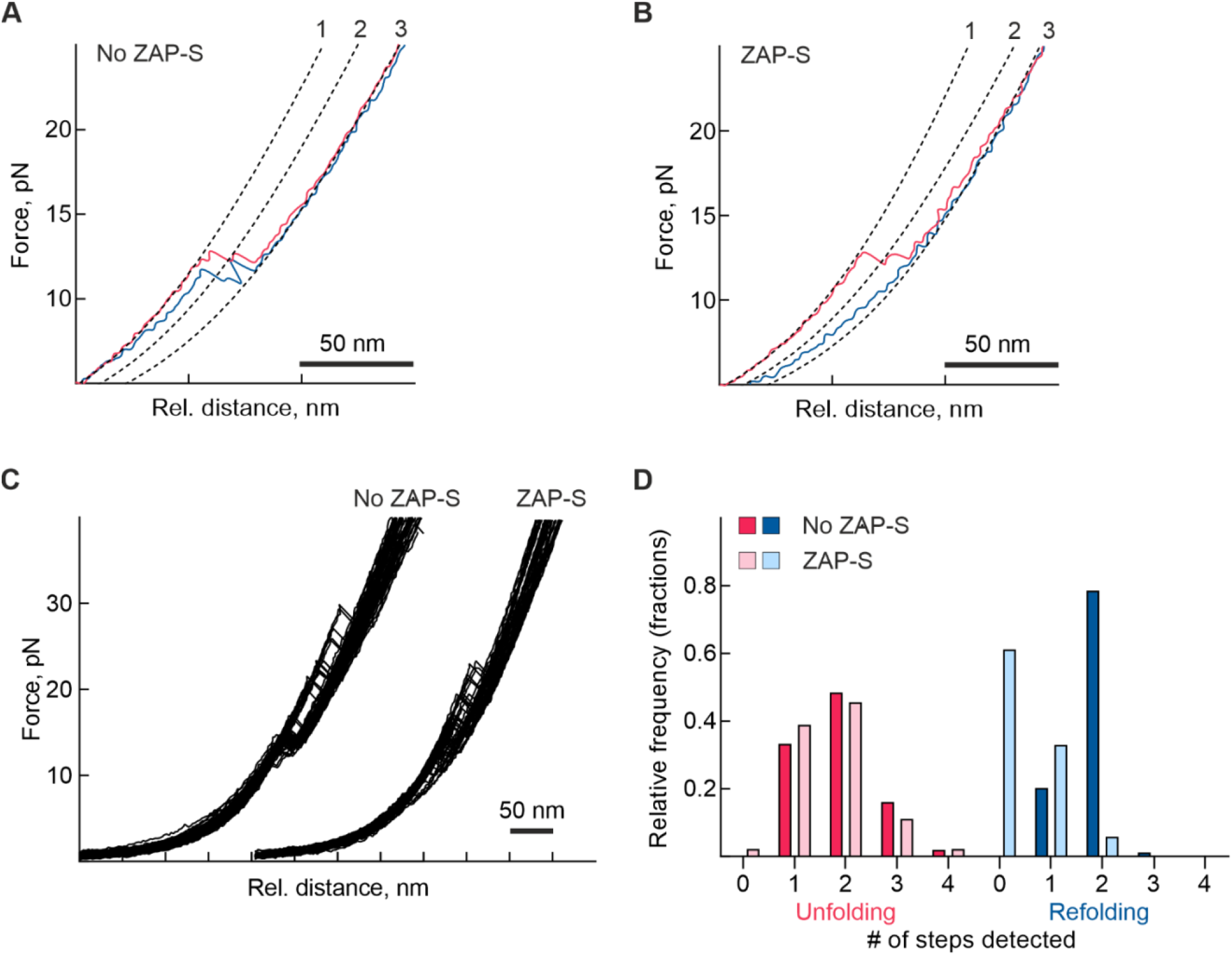
Optical tweezers data related to the Figure 5. **(A)** Example of the alternative unfolding (pink) and refolding (blue) force-distance curve of SARS-CoV-2 RNA sample (No ZAP-S). **(B)** Example of the alternative unfolding (pink) and refolding (blue) force-distance curve of SARS-CoV-2 RNA in presence of 400 nM ZAP-S (ZAP-S). Regardless the unfolding profile, the effect of ZAP-S remains. **(C)** Overlay of all force-distance curves used for the analysis of No ZAPS and ZAP-S samples. **(D)** A bar chart showing the number of unfolding (pink) and refolding (blue) steps per force-distance curve in RNA only (No ZAP-S, n=51) and RNA in the presence of 400 nM ZAP-S (ZAP-S) samples (n=45).

**Supplementary Fig. 5.**
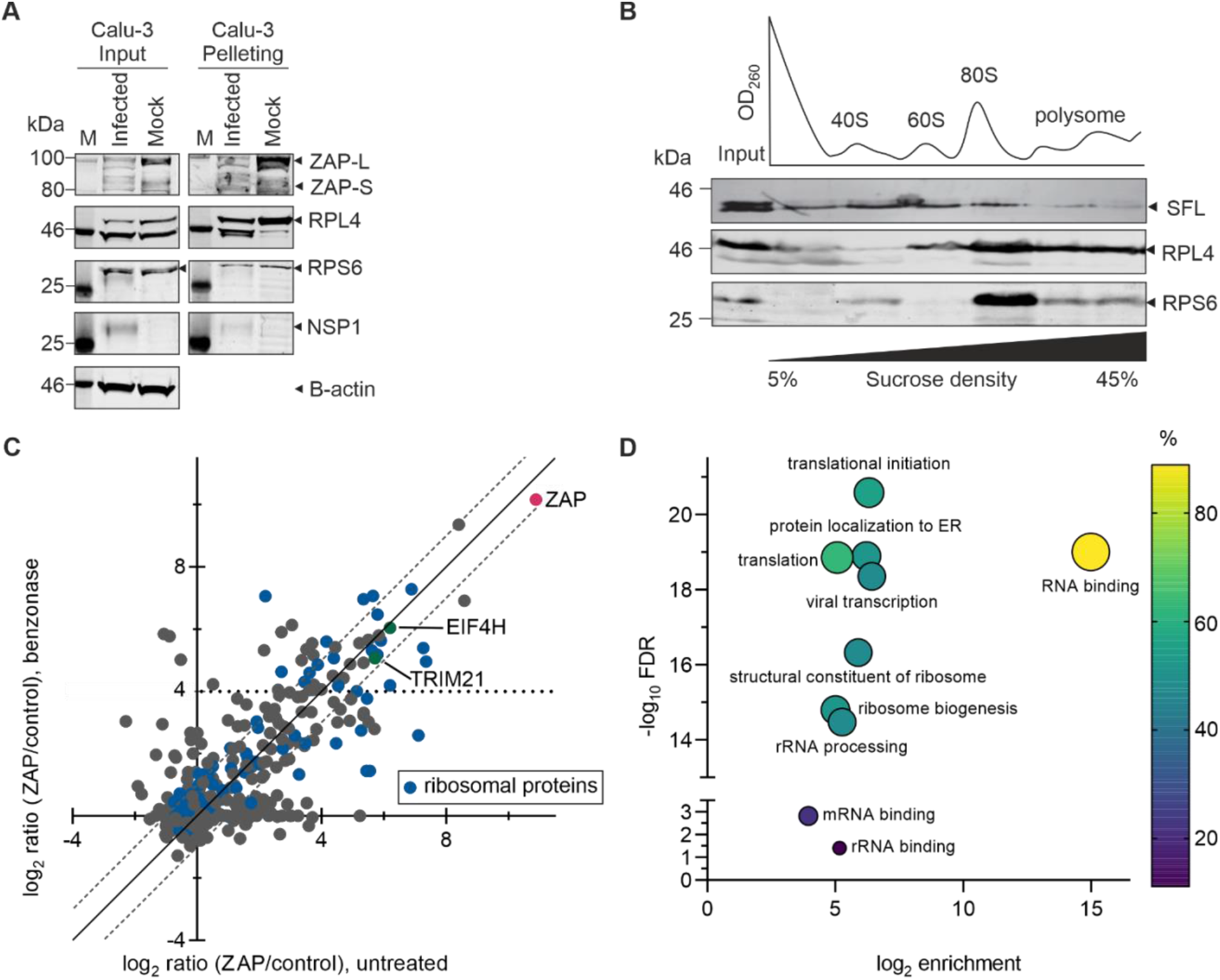
Interactions of *trans*-acting factors with ribosomes. **(A)** Ribosome pelleting of infected cells. Uninfected and SARS-CoV-2 infected Calu-3 cells were lysed and loaded onto 1M sucrose cushions. Levels of ribosomal proteins, ZAP, nsp1, and B-actin in the pellets were analyzed by western blotting using anti-RPL4, anti-RPS6, anti-ZC3HAV1 (ZAP), anti nsp1, and anti-beta actin antibodies. **(B)** Polysome profiling analysis for SFL protein. HEK293 cells transiently expressing SFL were lysed, subjected to 5-45% sucrose gradient ultracentrifugation, and subsequently fractionated. Levels of ribosomal proteins, as well as SFL in each fraction, were analyzed by western blotting using anti-RPL4, anti-RPS6, and anti-RYDEN (SFL) antibodies. **(C)** Scatter plot of the log_2_-transformed ratio of ZAP co-IP (ZAP-IP) with vs. without Benzonase. The dotted lines mark the difference in log_2_ ratio by 1 above and below the line of identity (solid line). **(D)** GO term analysis of proteins enriched by more than 16-fold in the ZAP-IP.

